# Two AGO proteins with transposon-derived sRNA cargo mark the germline in Arabidopsis

**DOI:** 10.1101/2022.01.25.477718

**Authors:** Gabriele Bradamante, Vu Hoang Nguyen, Marco Incarbone, Zohar Meir, Heinrich Bente, Mattia Donà, Nicole Lettner, Ortrun Mittelsten Scheid, Ruben Gutzat

**Author notes:** Swedish University of Agricultural Sciences, Uppsala, Sweden (Heinrich Bente). These authors contributed equally (Gabriele Bradamante, Vu Hoang Nguyen).

## Abstract

In sexually propagating organisms, genetic and epigenetic mutations are evolutionarily relevant only if they occur in the germline and provide inherited information to the next generation. In contrast to most animals, plants are thought to lack an early segregating germline, implying that somatic cells can contribute genetic information to the progeny. Here we demonstrate that two ARGONAUTE proteins, AGO5 and AGO9, mark an early-segregating germline. Both AGOs are loaded with dynamically changing populations of small RNAs derived from highly methylated, pericentromeric, long transposons. Sequencing single nuclei revealed that many of these transposons are co-expressed within an AGO5/9 expression domain of the shoot apical meristem (SAM). This indicates a host-parasite tug of war and specific silencing pathways along the plant germline throughout development. Our results open the path to investigate transposon biology and epigenome dynamics at cellular resolution in the SAM stem cell niche.

## 1. Introduction

All post-embryonic, above-ground organs of plants originate from stem cells in the center of the SAM which are marked by expression of *CLAVATA3* (*CLV3*) in Arabidopsis ^1^. Upon initiation of flowering, the vegetative SAM develops into an inflorescence meristem, which produces floral meristems. These develop floral organs, including stamens and carpels harboring male and female gametophytes. Both gametophytes develop within flower organ primordia from micro- and megaspore mother cells in the subepidermal (L2) layer ^2^.

Whether or not plants possess a separate group of germline cells prior to floral development and when germline identity is established is much debated ^3^. Early segregation would imply that germline cells should (i) be physically separated and hence recognizable by expression of specific genes, (ii) maintain a quiescent state with reduced cell cycle activity and high activity of DNA repair, and (iii), as predicted by evolutionary theory ^4^, display high expression levels of transposable elements (TEs) as well as genes regulating host defense mechanism. High expression levels of TEs and silencing-related genes have indeed been found in vegetative SAM stem cells of Arabidopsis ^5^ and rice ^6^, however addressing the question of germline identity in the SAM remains challenging, due to the difficulty of isolating and characterizing specific cell populations from shoot meristems.

ARGONAUTE (AGO) proteins are effectors in all small RNA (sRNA)-related pathways. The Arabidopsis genome contains 10 genes in three clades encoding AGO proteins. The AGO1/5/10 clade is associated with post-transcriptional gene silencing (PTGS) by binding to microRNAs (miRNAs) and targeting mRNA for degradation or translational inhibition ^7^. The AGO4/6/9 clade is associated with guiding sRNA-dependent DNA methylation (RdDM) to transposon sequences ^7^. RdDM activity can be recognized by DNA methylation in CHH (H demarks any base but G) context, mainly on short TEs on chromosome arms ^8^. Pericentromeric TEs are kept in a methylated heterochromatic state by the activity of the SWI/SNF2 chromatin remodeler DDM1 and the DNA methyltransferase CMT2 for establishing CHH methylation ^9,10^. In mutants lacking DDM1, transposons are massively transcribed ^11-15^, and binding of miRNA-loaded AGO1 to transposon transcripts triggers the synthesis of secondary siRNAs, thereby adding a PTGS layer to transposon repression ^16^. These 21/22 nt-long transposon-derived siRNAs, termed epigenetically activated siRNAs (easiRNAs) have also been found in male gametes ^17,18^.

We previously observed AGO5 and AGO9 expression in SAM stem cells ^5^ and hypothesized they might contribute to safeguard germline-precursor cells in the meristem from transposon invasion. Here, we characterized the spatial and temporal expression of AGO5 and AGO9 and their small RNA cargo. Their specific expression patterns in vegetative meristems allowed us to determine sRNA populations of L2 stem cells. To understand the role of both AGOs in TE repression, we employed transcriptome and methylome analysis of SAM stem cells in wild type and genetically perturbed plants. Taken together, we establish AGO5 and AGO9 as hallmarks of the Arabidopsis germline, which is established early in plant development, characterized by inflated TE expression and host counter defense, including the easiRNA pathway.

## 2. Results

### 2.1. AGO5 and AGO9 are present in germline and germline-precursor cells throughout development

To investigate the spatial distribution of AGO5 and AGO9 *in planta*, we generated reporter lines expressing both proteins with N-terminal GFP-tags under the control of their respective promoters and in the respective mutant background. *pAGO5::EGFP-AGO5* yields a specific signal in the cytoplasm of stem cells in the L2 of seedlings 7 days after germination (D7) (Fig. 1a; Supplementary Fig. 1). Throughout development, AGO5 also localizes to the L1 (Fig. 1c,e,w; Supplementary Fig. 2a,c) and is visible in axillary meristems (Supplementary Fig. 2i,k). During flower development, AGO5 is initially seen in the L1 of developing carpels (Fig. 1g,m; Supplementary Fig. 2e,g,m,q), male meiocytes (Fig. 1g,i; Supplementary Fig. 2m), and eventually in egg and sperm cells of mature gametophytes (Fig. 1k,o). During embryo development up to the octant stage, the AGO5 signal is uniformly distributed in the embryo proper (Fig. 1q; Supplementary Fig. 2s,u). In the globular stage, AGO5 appears restricted to the SAM L2, hypophysis and organizer (Fig. 1s), in the heart and torpedo stage to L2 and L3 of the SAM and to the root apical meristem (RAM) (Fig. 1u; Supplementary Fig. 2w), in agreement with ^19^. *pAGO9::EGFP-AGO9* is localized in nuclei mainly of the L2 in SAMs until floral induction (Fig. 1b,d, Supplementary Fig. 1). AGO9-labelled nuclei are also visible along the adaxial side of leaf petioles, apparently connecting to developing axillary meristems, where AGO9 is found at later time points (Supplementary Fig. 2b,j). In plants grown under a long day light regime (causing early flower induction), AGO9 is not found at D21 (Fig. 1f; Supplementary Fig. 2d,l) but is present in short-day-grown D21 plants, before flower induction, where it is restricted to the L2 (Fig. 1x, Supplementary Fig. 3). At the onset of flowering, the AGO9 signal seemingly migrates from the inflorescence meristem into floral meristems: initially seen between the whorls of carpels and stamens (Fig. 1h; Supplementary Fig. 2f,h), it is later found along the female and male lineages (Fig. 1j,n; Supplementary Fig. 2n,p,r). Like AGO5, it is present in egg and sperm cells of mature gametophytes (Fig. 1l,p) and in all nuclei of early embryos (Supplementary Fig. 2t,v). After the octant stage, it becomes gradually more restricted to the SAM region (Fig. 1t,v; Supplementary Fig. 2x), where it has been observed previously ^20^. These localization data are summarized schematically (Supplementary Fig. 4) and show that AGO9 is continuously present in nuclei of germ cells or their precursors throughout plant development. As the gametophytes develop from meristematic L2 cells, AGO9 labels the germline in plants according to the original Weismann definition ^21^. Although mostly cytoplasmic, AGO5 labels germ or meristematic L2 cells throughout most development in a pattern very similar to AGO9 (Supplementary Fig. 4). The cytoplasmic and nuclear preference for AGO5 and AGO9, respectively, suggest that both AGOs might act complementary for PTGS and TGS along the germline.

**Fig. 1:**
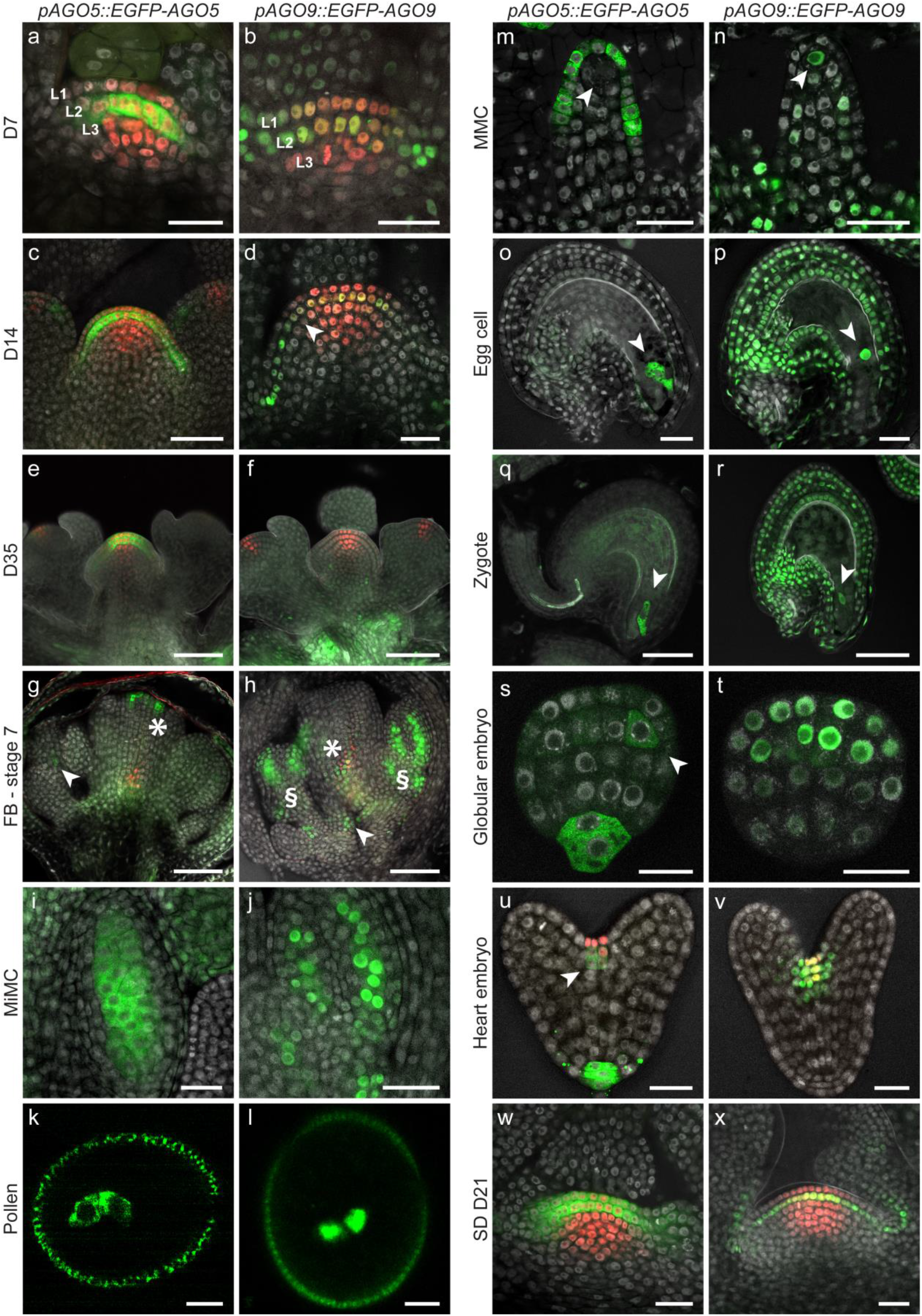
AGO5 and AGO9 are present in germline and germline-precursor cells throughout development. Expression of *pAGO5::EGFP-AGO5* and *pAGO9::EGFP-AGO9* (green) in lines containing the stem cell reporter *pCLV3::H2B-mCherry* (red) and lacking endogenous *AGO5* or *AGO9*. **a**, At D7 and **c**, at D14, AGO5 is localized mainly in the L2 layer of the apical dome of vegetative meristems. **b**, AGO9 is present in L2 and, to a lesser extent, in L1 at D7, whereas **d**, the AGO9 signal is restricted to L2 at D14. **e**, In inflorescence meristems at D35, AGO5 is uniformly localized in L1 and L2, whereas **f**, AGO9 is no longer visible. **g**, In stage 7 flower buds, AGO5 is localized in the upper region of the gynoecium (asterisk) and in the inner layers of future anthers (arrowhead). **h**, At the same stage, AGO9-labeled nuclei are found at the base of the bud (arrowhead) and in different cell layers in the upper developing organs, between the gynoecium (asterisk) and the inner layers of future stamens (section signs). **i**, In microspore mother cells, AGO5 is localized in the cytoplasm, whereas (**j**, a subset from 1h) AGO9 is detected in the nuclei. **k**, In mature pollen, AGO5 is restricted to the cytoplasm and (**l**) AGO9 to nuclei of sperm cells. **m**, In Stage 1 of ovule development, AGO5 is visible in the L1 of ovule primordia, whereas **n**, AGO9 is localized in the megaspore mother cell (arrowhead). **o**, With the formation of a complete embryo sac, AGO5 becomes restricted to the egg cell (arrowhead), whereas **p**, AGO9 is localized in nuclei of the egg cell and most of the surrounding sporophytic tissue. **q** + **r**, Zygotes (arrowhead) contain both AGO5 and AGO9. **s**, In the globular embryo, AGO5 is found predominantly in the L2 of the upper tier and the hypophysis and organizer cell, whereas **t**, AGO9 is present only in the upper tier. **u**, In the heart stage embryo, AGO5 is present in L2 and L3 of the SAM and in the root apical meristem (RAM). whereas (**v**) AGO9 is restricted to the SAM. **w**, In plants grown three weeks (D21) in short day light regimes, AGO5 is homogeneously localized in the two upper layers of the SAM, whereas **x**, AGO9 is restricted to the L2. Scale bars k, l = 5 µm; a, b, d, i, j, m, n, s, t, u, v, w, x = 20 µm; c, e, f, g, h, o, p, q, r = 50 µm.

Neither *ago5* nor *ago9* mutants have easily scorable morphological phenotypes, but *ago9* was reported to have an increased number of enlarged subepidermal cells in ovule primordia – the likely precursors of megaspore mother cells (MMCs) ^22^. To confirm the functionality of the tagged reporter lines, we asked whether they would complement this developmental defect. Unexpectedly, we could not observe the described difference between wild type and *ago9* mutant plants, possibly due to differences in growth conditions, as the number of enlarged subepidermal cells was relatively high also in the wild type. However, we detected a significantly increased number of enlarged subepidermal cells in *ago5 ago9* double mutants (Fig. 2). This phenotype could be rescued by introducing either *pAGO5::EGFP:AGO5* or *pAGO9::EGFP:AGO9* (Fig. 2), demonstrating that both tagged proteins are functional in the reporter lines. This further supports the hypothesis that AGO5 and AGO9 have partially redundant and complementary functions, in this case to restrict the number of MMC precursors in ovule primordia. We also wondered whether *ago5* and *ago9* could influence stem cell numbers. However, analysis of D7 SAMs of *ago5, ago9*, and *ago5 ago9* revealed no significant differences (Supplementary Fig. 5).

**Fig. 2:**
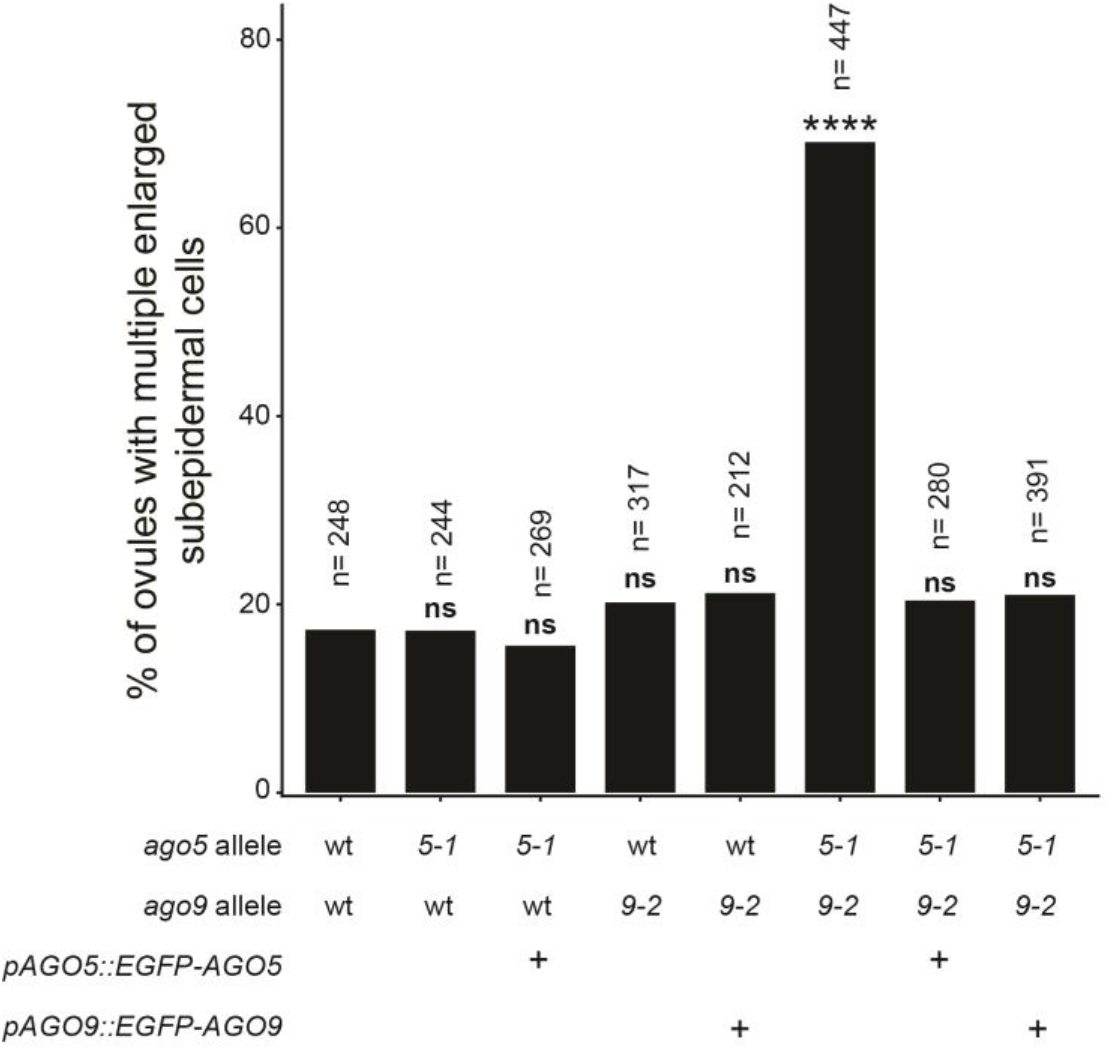
AGO5 and AGO9 have complementary functions to restrict the number of MMC precursors in ovule primordia. Gynoecia were analyzed by Nomarski interference contrast microscopy to determine the percentage of ovules with more than one enlarged subepidermal cell. Wild type and all mutant lines are in the *pCLV3::H2B-mCherry* background. Statistical significance by binomial test was performed against the wt. **** p-value < 0.0001, ns = not significant.

### 2.2. The sRNA cargo of AGO5 and AGO9 is dynamic and derived from transposons

To understand the function of AGO5 and AGO9 during Arabidopsis germline development and to assess sRNA pools from L2 SAM stem cells, we isolated and sequenced AGO5- and AGO9-bound sRNAs at two developmental time points (Fig. 3a, Supplementary Fig. 6,7). We chose shoot apices of D7 seedlings because of the specific AGO5 and AGO9 expression in L2 and L1/L2 (Fig. 1). To understand changes in AGO loading during germline differentiation, we also chose dissected apices from mature plants (D35) encompassing the inflorescence meristem, floral meristems, and very young flower buds. Protein levels of AGO5 are much lower than that of the prevalent AGO1 (Supplementary Fig. 7). To avoid contamination during D7 AGO5 precipitation, we introduced an AGO1 depletion step (Supplementary Fig. 7a). We confirmed a preferential AGO5 cargo consisting of 21, 22, and 24 nt sRNAs with a 5’ C bias (Fig. 3b, Supplementary Fig. 8), as was previously reported from cell cultures ^23^. AGO9 is loaded mainly with 5’ A-biased 24 nt sRNAs (Fig. 3b, Supplementary Fig. 8). Principal component analysis showed an increased variance of sRNA populations at D35 (Fig. 3c), suggesting diversification of sRNA populations in AGO5 and AGO9 during later development.

**Fig. 3:**
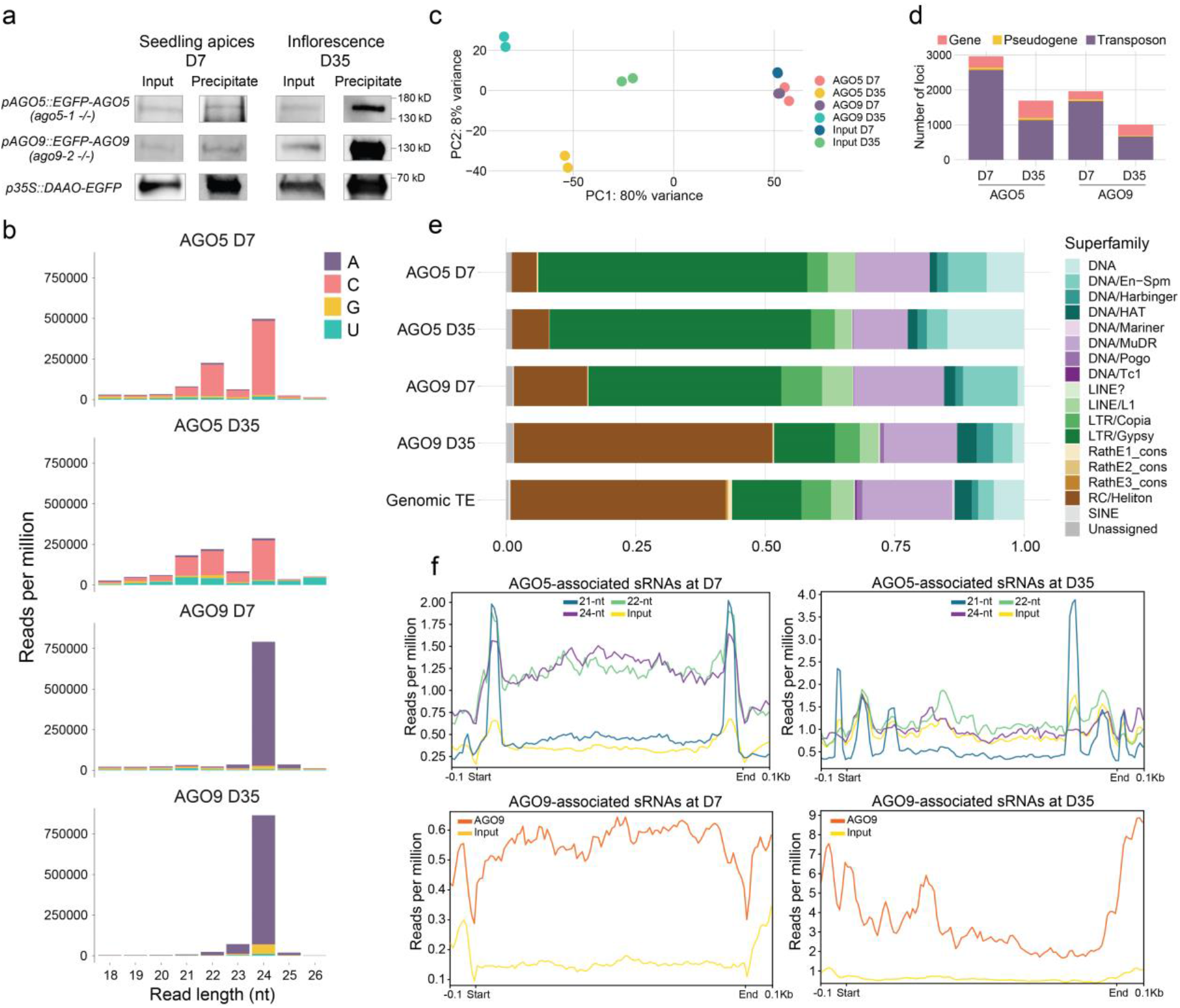
AGO5 and AGO9 sRNA cargo derived from transposons changes dynamically throughout development. **a**, Western blot of immunoprecipitation of GFP-tagged-AGO5 and AGO9 at D7 and D35. **b**, Read length distribution and 5’ bias of AGO5- and AGO9-associated sRNAs. **c**, Principal component analysis of the sequencing data. **d**, Number of target features of AGO5 and AGO9 (fdr < 0.05, Log2FoldChange >1). **e**, Potential transposon targets classified by superfamilies. **f**, Metaplots of sRNA distribution across the potential transposon targets.

A comparison between AGO-bound sRNAs with total sRNAs (input, FDR < 0.05, log2FoldChange >1) revealed that both AGOs contain mainly sRNAs homologous to transposons (Fig. 3d). AGO5 is predominantly associated with LTR/Gypsy retrotransposon-derived sRNAs, whereas AGO9 cargo contains proportionally more RC/Helitron sequences (Fig. 3e, Supplementary Fig. 9, Supplementary Table 1). The overlap between AGO5- and AGO9-targeted TEs is highly significant but less pronounced at D35 due to the strong bias towards Helitron-derived sequences in AGO9 (Fig. 3e, Supplementary Fig. 9). Notably, TEs represented in AGO5-sRNAs are pericentromeric, but AGO9-sRNAs targeted TEs shift from pericentromeres to chromosome arms during development (Supplementary Fig. 10).

AGO5-bound 21 nt sRNAs are mainly derived from LTR/Gypsy elements, similar to easiRNAs ^16^, and map most prominently to the 3’ and 5’ end of TEs, similar to profiles of easiRNAs in pollen ^17^. AGO5-associated 22/24 nt sRNAs are distributed more uniformly along TEs (Fig. 3f) as are AGO9-bound 24 nt sRNAs (Fig. 3f). The preferential loading of TE-related sRNAs implies that both, AGO5 and AGO9, are TE-silencing factors in the Arabidopsis germline throughout development.

### 2.3. *AGO5*- and *AGO9*-expressing cells show high expression of TEs

TE-derived siRNAs can be active either cell- or non-cell autonomously. A model for the male germline proposes the expression of TEs in companion cells and the migration of TE-derived siRNAs to gametes to reinforce RNA-directed silencing ^24-26^. We therefore wanted to understand whether this is similar in SAMs and asked whether the observed increase in TE expression ^5^ is confined to stem cells surrounding the L2 (analogous to “companion cells”), to L2 cells (analogous to future “gametes”), or uniform across all stem cells.

To test this, we FACS-sorted and analyzed transcriptomes of 188 individual *pCLV3::H2B-mCherry* nuclei derived from D7 plants using the SMART-seq platform (Supplementary Methods). We covered 21,055 genes and 3,706 TEs that were expressed in at least four nuclei (median of 3197 expressed genes and TEs per nucleus, Supplementary Fig. 20). To detect robust gene expression heterogeneity within this sparse dataset, we first adjusted for correlation between any two genes based on their total sampling ^27^ (Supplementary Methods). Expression patterns of three known cell cycle genes and 79 **g**enes **e**xpressed **s**pecifically in D7 **s**tem cells ^5^ (GESS) defined two major clusters, separating GESS into two groups (Fig. 4a, Supplementary Fig. 11). Besides *AGO5*, GESS group 2 comprises *MCT2* and *PHDG4*, two indicators for the L2 layer ^28^, and *CDT1A*, labeling cells in the G1 phase of the cell cycle (Fig. 4a). Noticeably, we found cluster 1 enriched for genes involved in the meiotic cell cycle, for silencing, and for microtubule-associated proteins, in agreement with the profile of L2 cells in the inflorescence meristems ^28^ (Fig. 4b,c). These data suggest that L2 cells of the SAM stem cell niche have a distinct expression pattern already early during vegetative development and are mainly in the G1 state of the cell cycle.

**Fig. 4:**
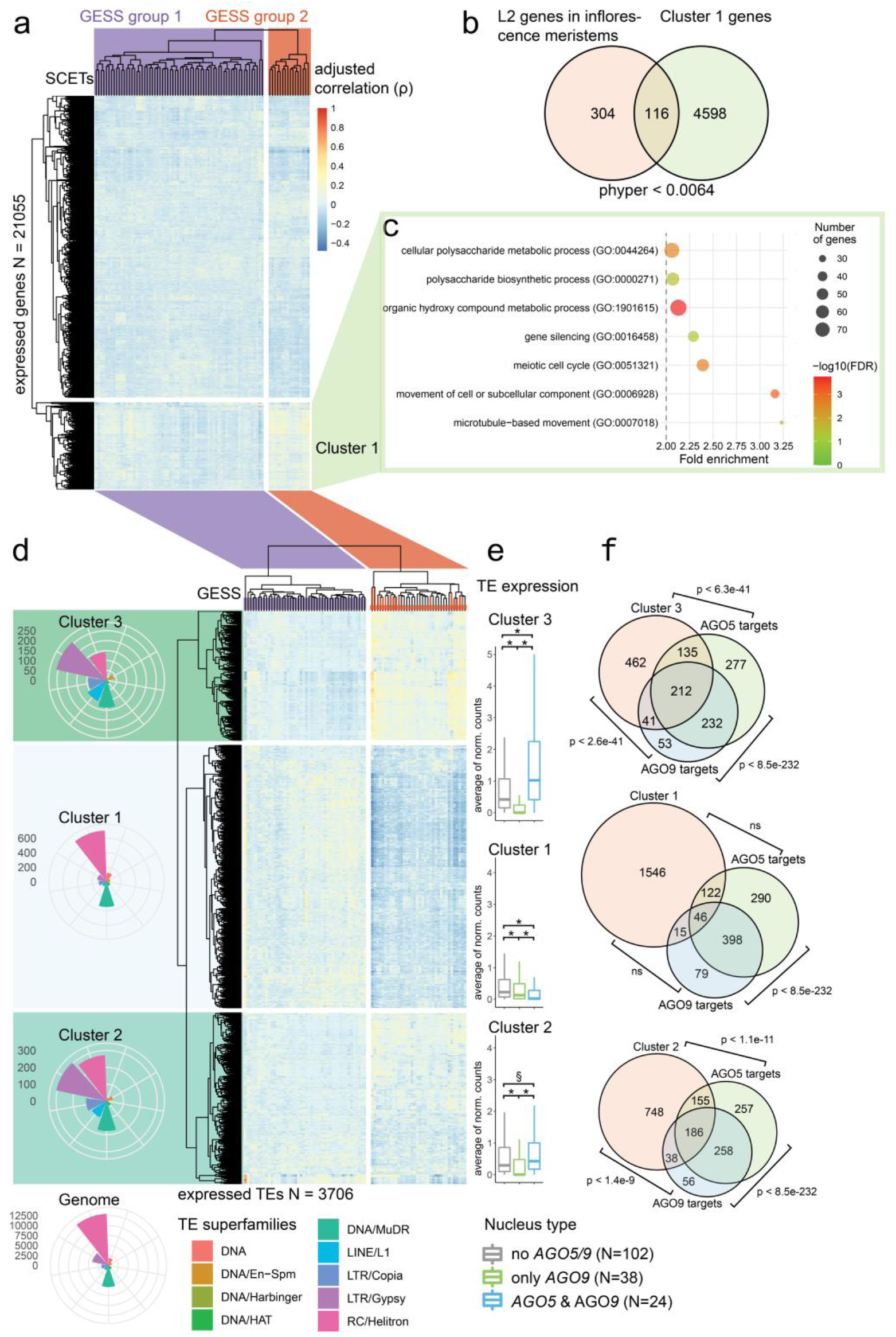
*AGO5*- and *AGO9*-expressing cells show high expression of TEs. **a**, Clustering of gene expression correlation with genes expressed specifically in D7 stem cells (GESS, see text) shows two major gene clusters.**b**, Cluster 1 genes are significantly enriched for L2 expressed genes according to ^28^. **c**, Most significant enriched GO-terms of genes in cluster 1. **d**, Clustering of TE expression correlation with GESS shows 3 major TE clusters. Group 1 GESS in the gene cluster also group together in the TE cluster (GESS group 2, red lines). Cluster 2 and 3 are highly enriched for LTR/Gypsy elements compared to the genome-wide distribution. **e**, Differential expression analysis shows high expression of TEs in cluster 2 and 3 in AGO5 and AGO9 containing nuclei. **f**, TEs with high expression in cluster 2 and 3 are highly enriched for AGO5 and AGO9 targets. (Venn diagram areas are not drawn proportionally). p = phyper, § = U-test < 1e-6, * = U-test < 2e-16

Performing the same correlation analysis between GESS and TE transcripts revealed three main TE clusters, separating two groups of GESS (Fig. 4d). All GESS from group 2 in the gene cluster were present in GESS group 2 in the TE cluster, suggesting that the TE expression pattern in GESS group 2 is mainly determined by L2 nuclei (Fig. 4d). Most TEs present in cluster 1 consist of RC/Helitrons and DNA/MuDR TEs, and their relative abundance resembles their genome-wide distribution (Fig. 4d), whereas TE clusters 2 and 3 are especially enriched for LTR/Gypsy transposons. Importantly, a highly significant portion of TEs from cluster 2 and 3 are targeted by AGO5-as well as AGO9-bound sRNAs (Fig. 4f).

To confirm this correlation analysis, we analyzed differentially expressed genes and TEs (DEGs) in nuclei grouped into those expressing both, AGO5 and AGO9, AGO9 only, or neither AGO5 nor AGO9. TEs from clusters 2 and 3 showed a significantly increased expression in “AGO5&AGO9” compared to “AGO9only” or “noAGO5/9” nuclei (Fig. 4e). Furthermore, AGO5- and AGO9-targeted TEs are also significantly longer than the genomic average of their respective superfamily (Supplementary Fig. 12). In summary, the data from single stem cell nuclei suggest the existence of at least two distinct niches of TE expression in stem cells, with high expression of LTR/Gypsy elements in AGO5-expressing cells, indicating cell-autonomous synthesis of TE-derived sRNAs. This raised the question whether this specificity is reflected in the specification of the DNA methylation state and transcriptional programs within the germline.

### 2.4. DNA methylation of heterochromatic TEs depends partially on AGO5

DNA methylation is highly dynamic in stem cells ^5^ and the male germline during differentiation ^29^, characterized by increased CHG methylation and decreased CHH methylation. Loss of CMT2 leads to reduced CHG and CHH methylation, especially on long heterochromatic TEs ^9^. Interestingly, AGO5- and AGO9-associated sRNAs are highly enriched for sequences matching TEs methylated by CMT2 rather than by the RdDM pathway (Fig. 5a) ^30-32^, suggesting reduced pericentromeric methylation and hence TE expression in cells expressing AGO5 and AGO9. To test this, we performed DNA methylation analysis on stem cells and male germ cells by sorting and collecting D7 stem cells and sperm nuclei of wt, *ago5, ago9*, and *ago5 ago9*. We observed a slight reduction of CHG and CHH methylation in stem cells of *ago5* and *ago5 ago9* seedlings, and this reduction was significantly more pronounced in sperm cells (Fig. 5b). CHG and CHH methylation in sperm cells of *ago5* and *ago5 ago9* was especially reduced on TEs longer than 1500 bp (Fig. 5c). Total methylation levels of TEs matching AGO5 and AGO9 cargo were significantly higher than at non-targeted TEs (Supplementary Fig. 13), and reduction of CHG methylation was most prominent in sperm in the double mutant *ago5 ago9* at TEs that corresponded to the most abundant AGO5/9-bound sRNAs (Fig. 5d). Unexpectedly, most of the reduction in CHH and CHG methylation at TEs was correlated with the lack of *AGO5*, as *ago9* had only minimal additional effects in the double mutant (Fig. 5b). This influence of AGO5 on DNA methylation could either be indirect, by repressing the mRNA of genes in the cytoplasm, or direct by shuttling of AGO5 with its cargo into the nucleus. Nuclear shuttling has been demonstrated for AGO1 ^33,34^, and AGO1 and AGO5 share high sequence similarity at the N-terminus, including a nuclear export signal (NES). We mutated the AGO5 NES and observed a significant accumulation of nuclear AGO5-NESm (Supplementary Fig. 14). This, together with the 24 nt sRNAs bound to AGO5, suggests that AGO5 could have a role in both, cytoplasm and nucleus, and may play a role in PTGS as well as TGS linked with DNA methylation.

**Fig. 5:**
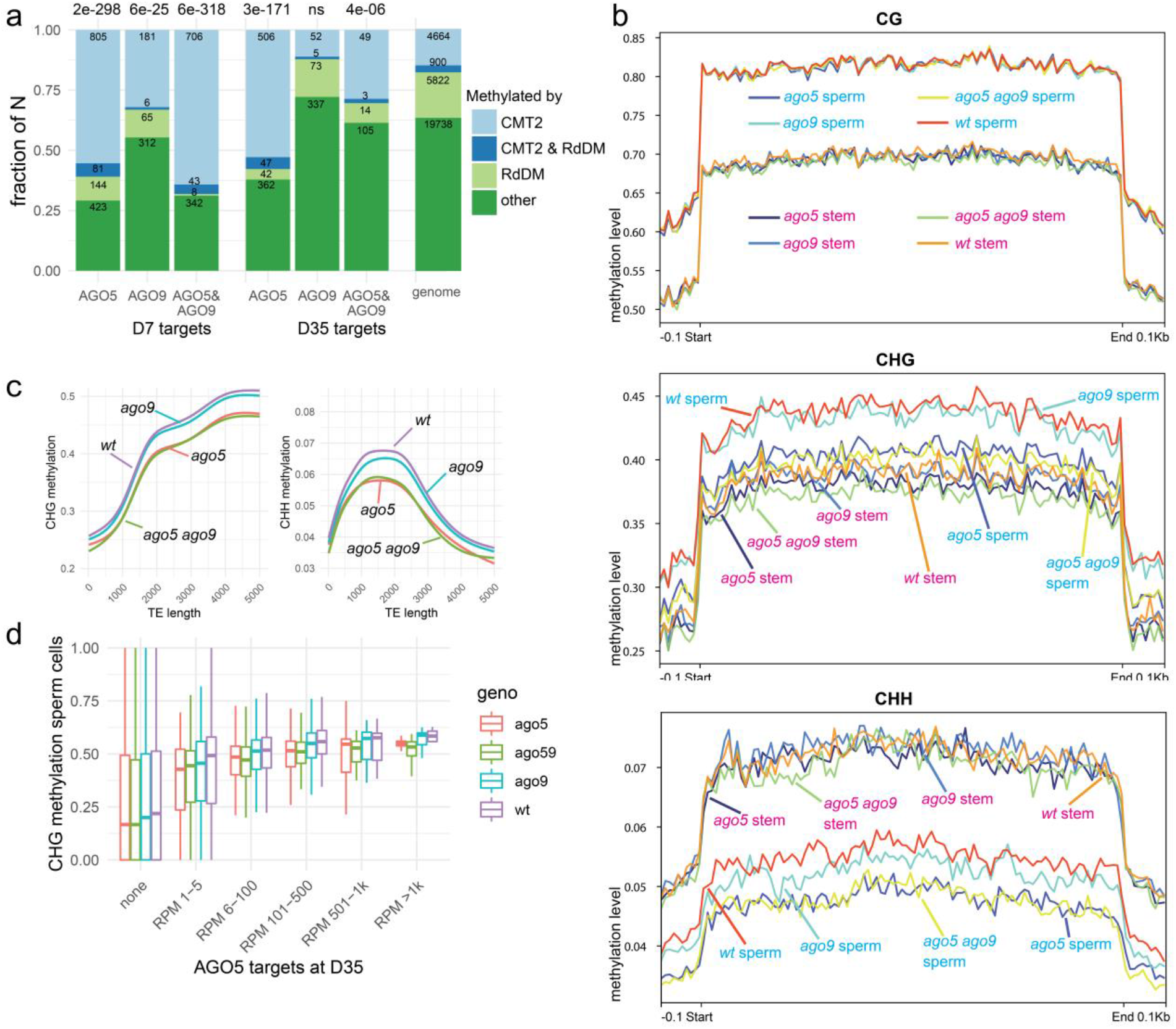
DNA methylation of heterochromatic TEs depends partially on AGO5. **a**, AGO5 and AGO9 targets are mostly methylated by CMT2 in seedlings. Numbers above bar plots indicate p-values (phyper) for the enrichment of CMT2 targets. Numbers in the bar plots indicate the number of respective targets. **b**, CG, CHG, and CHH methylation on TEs. CHG and CHH methylation is reduced in sperm cells of *ago5* and *ago5 ago9*. **c**, TE methylation level over TE length. **d**, CHG methylation levels in sperm cells of TEs targeted by AGO5 and sorted by the abundance (RPM) of AGO5-associated sRNAs.

### 2.5. TEs corresponding to AGO5 and AGO9 cargo are derepressed when DNA methylation is impaired

To address whether the loss of AGO5 and AGO9 results in increased transcription of the TEs corresponding to their cargo, we sequenced mRNA of D7 stem cells and non-stem cells of the SAM, and sperm and vegetative nuclei of pollen, of wt, *ago5, ago9*, and *ago5 ago9*. Expression of *CLV3, mCherry, DUO1, MGH3*, and *VCK1* demonstrated enrichment of the respective cell types (Supplementary Fig. 15b). Several *AGO* genes are highly expressed in D7 stem cells, while only *AGO5* and *AGO9* transcripts are detectable in sperm cell nuclei (Supplementary Fig. 15c). This suggests non-redundant functions of these two AGOs in sperm cells. However, the nuclear transcriptome of sperm and vegetative cells differed only minimally between the different genotypes, and only 6 TEs showed increased expression in *ago5 ago9* sperm nuclei (Supplementary Fig. 15a), a surprising result as *AGO5* and *AGO9* were the only *AGO* genes expressed in pollen.

Among the AGO4/6/9 clade of AGO proteins, AGO4 is crucial for TGS in seedlings ^31^ and is highly expressed in stem cells (Supplementary Fig. 15c). To probe for redundancy of AGO9 with AGO4 in vegetative meristems, we created additional double and triple mutants and sequenced mRNA from shoot apices of D7 seedlings of wt, *ago4, ago5, ago9, ago4 ago9, ago5 ago9*, and *ago4 ago5 ago9*. To also probe for a potential connection to the easiRNA pathway, we included *ddm1. ddm1* is characterized by a global loss of DNA methylation, strong de-repression of long and heterochromatic transposons, and emergence of easiRNAs ^16^.

Comparisons between a*go* mutant shoot apex transcriptomes revealed only a few differentially expressed genes and TEs (Supplementary Fig. 16). As expected, more than 1300 TEs were derepressed in *ddm1* compared to wt. Interestingly, these TEs displayed a significant overlap to those specified by the AGO5 and AGO9 cargo (Fig. 6a), and overlapping TEs were more highly expressed in *ddm1* than those not represented among AGO5-/AGO9-associated small RNAs (Fig. 6b). Hence, TEs that are precursors of AGO5- and AGO9-associated sRNAs – and potentially targeted by these AGOs – react strongest to the loss of DNA methylation.

**Fig. 6:**
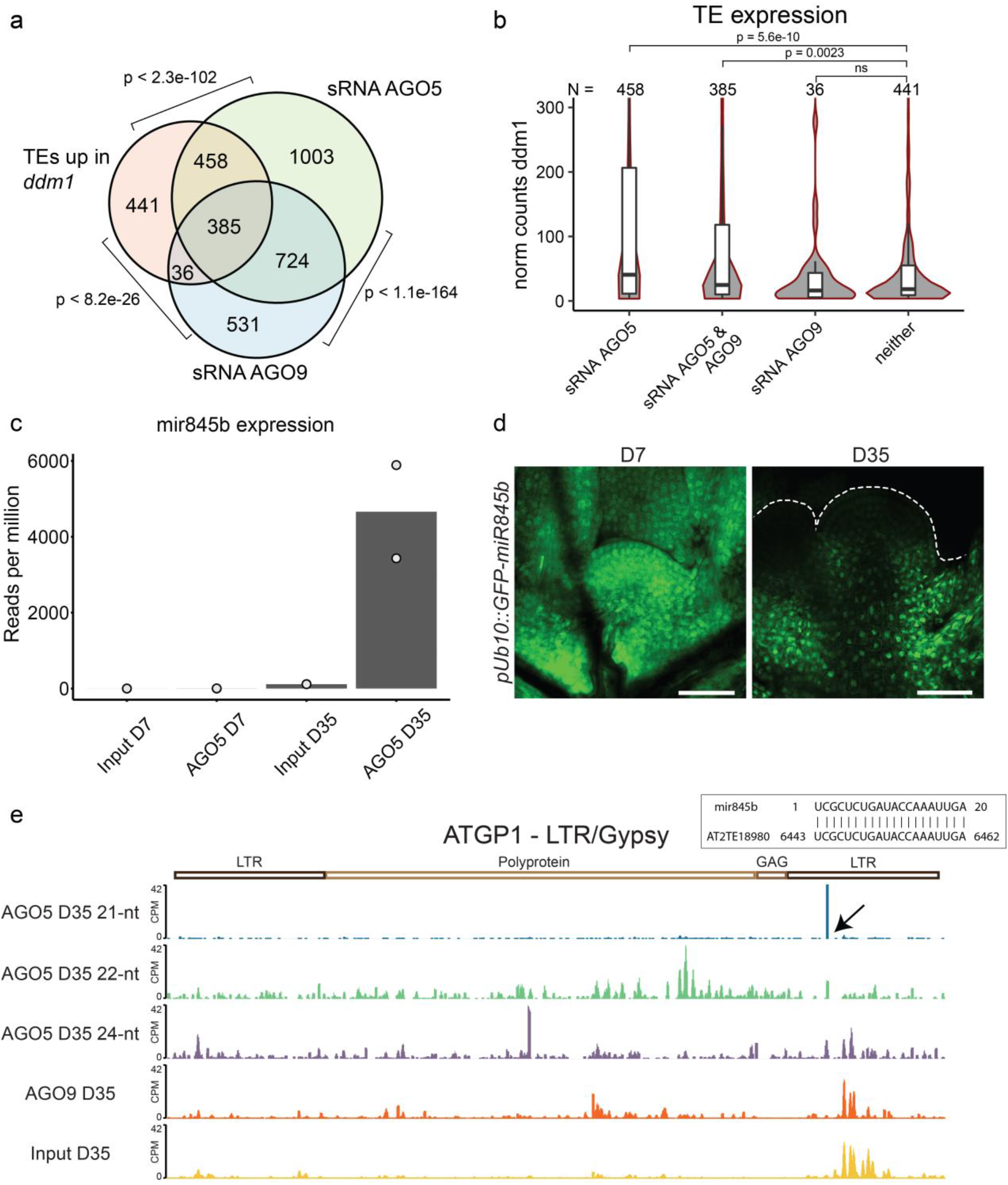
AGO5 is an easiRNA pathway effector. **a**+**b**, AGO5 and AGO9 cargo is derived from TEs with high expression in *ddm1*. **c**, mir845b association with AGO5 at D7 and D35. **d**, *pUb10::GFP-miR845b* reports on mir845 activity. Scale bar b = 50 µm. **e**, Integrated genomic viewer of an LTR/Gypsy element, the arrow indicates target site of miR845b.

These data show that *ddm1*-sensitive, long pericentromeric TEs are expressed in AGO5- and AGO9-containing stem cells, and siRNAs derived from these TEs are incorporated into AGO5 and AGO9. Subsequently, cells expressing AGO5 must allow either PolII or PolIV access to pericenteromeres to generate the precursors. easiRNA synthesis is initiated by the activity of miRNAs, and we found miRNA845, a crucial trigger of easiRNA biosynthesis ^18^ and other potentially TE targeting miRNAs, significantly associated with AGO5 in SAM stem cells (Fig. 6c and Supplementary Table 2). To test whether mir845 could be involved in silencing TEs in stem cells, we used the UBQ10p::GFP-miR845b reporter, a sensor of mir845b activity that results in loss of the GFP signal ^18^. GFP fluorescence is visible in stem cells at D7, probably due to the low abundance of mir845b (Fig. 6c,d), but it is strongly reduced in inflorescence meristems at D35, congruent with high levels of mir845b associated with AGO5 (Fig. 6c). An example of a transposon potentially targeted by mir845b is shown in Fig. 6e.

Synthesis of easiRNAs in *ddm1* depends on RDR6, but in pollen seems to depend on POLIV ^16,17^. Therefore, we wanted to understand the synthesis of TE-derived sRNAs in shoot apices and performed northern blot analysis to probe for AGO5-targeted TEs in mutants involved in sRNA generation. This revealed a dependency on DCL1-3, POLIV, and RDR2 (Supplementary Fig. 17), similarly to the situation in pollen and embryos ^32^. Taken together, our results imply that AGO5 is a native easiRNA effector in SAM stem cells.

## 3. Discussion

Our study presents strong evidence for an early segregating germline with a specific complement of epigenetic effectors in Arabidopsis. Despite the small number of SAM stem cells, the expression patterns of AGO5 and AGO9 demonstrate heterogeneity among them. Co-expression analysis of single stem cell nuclei revealed two stem cell niches displaying increased expression of transposons in *AGO5-* and *AGO9-*expressing cells. sRNAs derived from these TEs are loaded into AGO5 and AGO9, and likely help prevent TE mobilization by reinforcing DNA methylation at CHG and CHH sites, as well as by PTGS. Interestingly, AGO5 also binds to miRNA845, a crucial trigger of easiRNA biogenesis necessary for silencing hundreds of LTR/Copia and LTR/Gypsy elements in *ddm1* ^16^.

Moreover, we found that a large portion of AGO5- and AGO9-targeted TEs are expressed in *ddm1*, although we observed increased levels of *DDM1* transcripts in bulk stem cells ^5^. This suggests a chromatin state permissive for TE expression in AGO5- and AGO9-containing stem cells. Alternatively, populations of TEs could be activated by stem cell-specific transcription factors or signaling networks. In contrast to AGO1, which has a high affinity for 5’U-containing sRNAs ^23^, AGO5 has a bias for 5’C sRNAs, which probably prevents competition with AGO1 for sRNA duplexes. This suggests a functional specialization of AGO5 for post-transcriptional TE silencing via easiRNAs in the Arabidopsis germline. However, we do not find strong de-repression of TEs in the absence of AGO5 or AGO9, likely because AGO1 is still able to trigger easiRNA biosynthesis.

Interestingly, several plant clades show AGO1/5/10 diversification ^35^, and the AGO5 homologs in maize, MAGO1 and MAGO2, are crucial for preventing TE mobilization during male gametogenesis during heat stress ^36^. Unexpectedly, we found only *AGO5* and *AGO9* expressed in pollen, although AGO1 can silence the mir845-reporter ^18^. Carry-over of other AGO proteins from the microspore could explain this observation. The role of both AGOs in female gametogenesis needs to be further investigated, especially since LTR/Gypsy elements seem to be also expressed in egg cells ^37^. AGO5 was reported to be involved in megasporogenesis; however, this result was obtained with a truncated, dominant allele of AGO5 lacking the ability to selectively bind sRNAs ^19^. Polymorphisms in natural Arabidopsis accessions in the *AGO9* gene are associated with DNA methylation variation across TEs ^30^, but we could find only minor methylation differences in *ago9* stem or sperm cells. However, transposons fixed in the population are very old and have likely lost their ability to mobilize. If AGO9 is necessary for *de novo* DNA methylation, or if it contributes indirectly to variation of DNA methylation by post-transcriptional silencing, an effect on DNA methylation in *ago9* mutants might not be detectable.

How far the results from Arabidopsis reflect the situation in other plants needs further studies, but they indicate remarkable similarities with germline stem cells in animals. For example, deleting piRNA pathway components leads to strong activation of TEs in gametes and gamete-companion cells in the gonads of Drosophila, allowing different TE families to mobilize with varying strategies ^38^. While Arabidopsis, and plants in general, have diverse and partially redundant TE silencing pathways, studying gene and TE expression in single stem cells at different developmental stages, combined with information about the (sub-)cellular localization of the proteins in wild type and mutants, will provide unprecedented insight into the complex interplay of transposon mobility and silencing along the germline.

## 4. Material and Methods

### Plant material

Experiments were performed with *Arabidopsis thaliana* ecotype Col-0. Mutant and reporter lines used are listed in Supplementary Table 3. AGO5 and AGO9 reporters were cloned into pElvis; pElvis was generated from pSun ^39^ by inserting an additional marker conferring seed fluorescence: pSun was linearized using EcoRV; next, a functional OLE1::GFP expression cassette ^40^ was assembled from two PCR fragments containing promoter::CDS and GFP::terminator (fragment from pEarlyGate103 ^41^) respectively, and inserted into pSun using In-Fusion^®^ cloning (Takara Bio Cat. #121416) according to manufacturers’ instructions.

pAGO5::EGFP-AGO5 was constructed amplifying a ∼6 kb genomic fragment containing the ORF and ∼500 bp 3’ sequence and inserting it into pElvis using HindIII and PmeI sites. Next, a ∼2.5 kb promoter fragment was inserted via KpnI and HindIII sites. Finally, EGFP was inserted using HindIII and In-Fusion^®^ cloning (Takara Bio Cat. #121416). For pAGO9::EGFP-AGO9, a ∼5 kb fragment containing the ORF and ∼500 bp 3’ sequence was inserted into pELVIS using Kpn1 and BamH1. Next, the vector was cut with Kpn1, and a ∼3 kb promoter fragment containing the 5’ UTR of AGO9 was inserted. A Kpn1 site remained, and EGFP was inserted using In-Fusion^®^ cloning (Takara Bio Cat. #121416). For p35S::DAAO-GFP, a fragment containing the CaMV35S promoter and DAAO-GFP was blunt inserted into pSUN using SmaI and HindIII filled up with Klenow fragment.

pAGO5::Clo-AGO5NESm was engineered using the GreenGate system ^42^ by assembling the pGGA-pAGO5, pGGB-Clover, pGGC-AGO5NESm, pGGD-D-dummy, pGGE-3UTR-AGO5, and pGGF-YFP-seed-coat entry modules into pGGSun (pSun adapted for the Greengate system). For the CRISPR *ago4* lines (*ago4-CR)*, sgRNAs were designed *in silico* using “CHOPCHOP” ^43^. Three sgRNAs were chosen and tested with *in vitro* cleavage assay as described ^44^. sgRNAs that showed good cleavage efficiency on PCR products were cloned into a modified version of pDE-Cas9 ^45^, as described earlier ^46^ by using the tRNA multiplex system ^47^ and two pre-annealed oligonucleotides for each sgRNA. The resulting sgRNA cassettes were amplified with primers containing appropriate restrictions sites (MluI) and cloned into pDEECO vector ^44^. The two selected sgRNAs were matching against the first exon as well as the first intron of the *AGO4* gene (At2g27040). Plants were genotyped for a approx. 100 bp deletion in exon one including the start codon.

All oligonucleotides that were used in the study are described in Supplementary Table 3. Plants were transformed with the floral dip method, and transgenic seeds were selected under a fluorescence binocular, based on the expression of the oleosin-GFP encoded in the plasmid backbone.

### Growth conditions

All plants were grown either *in vitro* on GM medium with or without selection or on soil under 16/8 h or 8/16 h light/dark cycles (for long and short-day regimes, respectively) at 21°C with 60% humidity and 150 µmol m^−2^ s^−1^ light intensity. Plant material was always harvested at the same time of the light period. All plant lines and transgenic lines produced are described in Supplementary Table 3.

### Fixing and clearing of plant tissue

All plant tissue except mature pollen was fixed and cleared prior to microscopy according to the following procedure. Samples were first fixed in a 2% FAA solution as described in ^48^, for 10 min under vacuum and then placed on a thermoblock for 40 min at 37°C. Fixative was removed, and samples were incubated in ClearSee solution ^49^ at 4°C for two to seven days. Seven-day-old seedlings were incubated in ClearSee for four days; for older plants, leaves were first removed, and the remaining shoot was fixed and incubated for seven days. For inflorescence meristems, shoot tips from 35 day-old plants were placed on a Petri dish half-filled with 2% agarose, covered with distilled water, and dissected with a needle attached to a syringe to expose the SAM. Explants were fixed, cleared for two days, and the main stem removed prior to slide preparation. Gynoecia for the observation of egg cells and very young embryos were fixed and cleared for seven days. Ovules with globular, heart-stage, and torpedo embryos were collected from siliques and observed after fixing and clearing for seven days.

One day prior to microscopy, samples were stained with 1 mg/mL DAPI in ClearSee, except for gynoecia and ovules that were stained for the whole week of clearing. Samples were washed and mounted on Superfrost microscope slides with ClearSee. Mature pollen was released by vortexing detached flowers in a 0.3 M mannitol solution.

The pollen suspension was pelleted by centrifugation for 1 min and resuspended in 20 μl of the same solution. The whole suspension was loaded onto a Superfrost microscope slide for microscopy. Microscopic analysis was performed with an LSM880 Axio Observer with Airyscan detector.

### Microscopy of enlarged subepidermal cells in ovule primordia

Gynoecia at different developmental stages were dissected with forceps and scalpel and fixed overnight in 4% FAA (4% formaldehyde, 5% acetic acid, 50% ethanol), then dehydrated in 70% ethanol, cleared in Herr’s solution and observed on a Zeiss Axioobserver Z1 with differential contrast optics.

### Counting of stem cells

Images of meristems of seven day-old plants expressing H2B-mCherry from the CLV3 promoter were acquired as 16-bit z-stacks with the same settings for all genotypes examined. Segmentation and counting of H2B-mCherry-labelled stem cell nuclei were computed with Imaris 9.5.0 software. Nuclei were identified as single spots, and segmentation parameters were set to recognize spots only in the core of stem cell nuclei. The same parameters were applied for all acquisitions: Spots; Points Creation Parameters, Estimated Diameter: 3.250 3.250 3.250; Background subtraction: selected, Filter Type: quality; Lower Threshold Manual Value: 247, Upper Threshold Manual Value: 1.

### Quantification of cytoplasmic versus nuclear GFP

The ratio of cytoplasmic to nuclear GFP signal intensity was quantified in meristems of 35 day-old plants after acquiring 16-bit images with the same settings in the GFP channel for each line. Cell selection for segmentation was performed based on clarity of cell features and non-overlap with adjacent cells. Perimeter segmentation of the cytoplasm and the nucleus was manually drawn in Fiji for each cell, and the watershed function was applied to smooth edges. The average GFP intensity signal for the cytoplasm and nucleus area was then calculated. The ratio of cytoplasmic to nuclear GFP intensity for each meristem represents the average value of the selected cells. Steps were automatized by using a dedicated Fiji macro.

### Fluorescence-activated nuclei sorting (FANS)

The sorting of stem cells is described in detail ^50^. Pollen was harvested from flowering Arabidopsis plants as described ^51^. A vacuum cleaner was equipped with 150 µm and 60 µm filter meshes to block unwanted plant material and debris, and pollen was collected on a final 10 µm mesh. The pollen was transferred to Eppendorf tubes and stored at -80°C in aliquots of ca. 20 µL. The pollen was resuspended in 500 µL of Galbraith buffer and processed as described ^52^ to release sperm and vegetative nuclei. The nuclei were stained by adding 0.5% v/v SYBR-Green. The resulting suspension was directly subjected to FANS. Sperm and vegetative nuclei were sorted on a BD Aria III cell sorter (70 µm nozzle). A 488 nm Blue Laser, Coherent Sapphire 20 mW, was used to excite SYBR-Green, and signals were detected with a FITC 530/30 nm bandpass filter. Sorting gates were adjusted according to the different emission intensities between sperm and vegetative nuclei populations. DNA and RNA isolation was performed as described ^50^.

### AGO5- and AGO9 immunoprecipitation and sRNA preparation

Meristems of D7 and D35 plants transgenic for GFP-tagged AGO5 and AGO9 in the background of respective mutants were manually collected on ice. Material from 600 plants (D7) and 200 mg (D35) was frozen and ground in liquid nitrogen and powder mixed with IP buffer (20 mM, HEPES pH 7.5, 100 mM KCl, 0.2% NP-40, 10% glycerol, 1 mM EDTA, 1 mM PMSF, 20 µM MG132, 5 mM DTT and Roche protease inhibitor #5892953001), and incubated for 1 h on a rotating wheel. This and all future steps were performed at 4°C. Cell debris was removed by centrifuging twice for 10 min at 12000 g. Next, supernatants were precleared by incubation for 1 h with 200 µL control beads (Chromotek #bmab-20). For the 7 day-old meristem samples, an additional step was applied to deplete AGO1 by adding prepared 10 µL anti-AGO1 (Agrisera #AS09 527) with 50 µL beads (Invitrogen #10001D) and incubated for 30 min. This step was repeated once more. After bead removal, the supernatants were incubated with GFP-trap beads (Chromotek #gtma-10), 5 µL for the 7-day sample and 20 µL for the 35-day sample and incubated on a rotating wheel for 1 h. The beads were washed 5 times with immune-precipitation buffer. A third of the precipitate was used for western blots, and two-thirds were processed for RNA extraction in TRIzol (Invitrogen #10296010) reagent.

For western blots, the precipitation was mixed with 20 µL Laemmli buffer and incubated for 10 min at 95°C. After removing the beads, the mix was loaded on mini-PROTEAN stain-free gels (Biorad #4568083). Gel electrophoresis was done for 90 min at 30 mA, and the gel was washed in transfer buffer (20% methanol, 0.4% SDS, 48 mM Tris, 39 mM glycine). Protein was transferred to a nitrocellulose membrane (Biorad #162-0113) by semi-dry electroblotting (Biorad) at 20 V for 90 min. The membrane was incubated with anti-GFP primary-(Roche #11814460001/1:2000) and anti-mouse (Cell Signaling Technology #7076/1:5000) secondary antibody. The membrane was washed 3 times before image acquisition by ChemiDoc Touch Imaging System (Biorad). sRNA libraries were constructed with QIAgen miRNA library (QIAgen #331502) and were sequenced on an Illumina HiSeqV4 SR50. All steps were performed by the Next Generation Sequencing Facility (Vienna BioCenter Core Facilities).

### Small RNA data analysis

Raw reads from sRNA library sequencing were trimmed using cutadapt v1.18 ^53^, and 18 to 26 nt long reads were selected. The reads were aligned to the Arabidopsis genome (TAIR10 plus TAIR8 transposons as described in Supplementary Methods) using bowtie2 v2.3.5 ^54^, allowing 1000 times multi-mapping. The 5’ nucleotides of 18-26 nt sRNAs were analyzed using a pipeline available on GitHub (https://github.com/AlexSaraz1/paramut_bot). Subsequent data analysis was performed with 21 to 24 nt long reads. Counting of reads was done using featureCounts from the Subread package v2.0.1 ^55^. Differential expression analysis was performed by DESeq2 v1.32 ^56^ (fdr < 0.05, log2FoldChange > I1I). Genomic features having less than 5 normalized reads were filtered out. Deeptools v.3.3.1 ^57^ was employed to generate normalized count bigwig files using “bamCoverage” with the “CPM” parameter. Bigwig files merged from both replicates were used to generate metaplots. Blast+ ^58^ was employed to find potential targets of AGO5-bound miRNAs. Genomic TE sequences were blasted with the parameter “-task “blastn-short”” for short sequences and with miRNAs as input.

### Library preparation and sequencing

For single nuclei, RNAseq (snRNAseq) nuclei of shoot apices of 7 d old seedlings were prepared according to ^50^. Single nuclei were sorted into 96-well plates provided with 4 ul smart-seq lysis buffer ^59^. Library preparation and sequencing were performed by the Next Generation Sequencing Facility (Vienna BioCenter Core Facilities). For mRNA seq of sorted stem and non-stem nuclei, bulks of 100 nuclei were sorted into 96-well plates and proceeded as with single nuclei. For mRNA seq of sperm and vegetative nuclei and D7 shoot apices, total RNA was extracted. Smart-seq2 and 3 sequencing libraries and subsequent sequencing was performed by the Next Generation Sequencing Facility (Vienna BioCenter Core Facilities). For bisulfite library preparation, libraries were prepared with the Pico Methyl-Seq Library Prep Kit (Zymo Research #D5456) and sequenced by the next Generation Sequencing Facility (Vienna BioCenter Core Facilities). Sequencing information is compiled in Supplementary Table 4.

### Analysis of sequencing data

mRNA sequencing reads were processed with nf-core/rnaseq ^60^. Due to the redundancy of the TAIR annotations “transposable element” and “transposable element gene,” we used a custom annotation file containing TAIR10 features plus “transposable elements” without “transposable element genes” and added sequences of transgenes (see Supplementary Methods for details). Differential gene expression analysis was performed with DESeq2 ^56^. GO enrichments were calculated using the AmiGO2 tool and the PANTHER classification system (http://amigo.geneontology.org/rte) ^61^. Bisulfite sequencing data were processed with nf-core/methylseq ^62^. Visualization of the data was achieved using R and Bioconductor ^63^ including the packages tidyverse, ggplot2, pheatmap and a protocol for GO-term enrichment analysis ^64^.

### Northern blotting

Twelve μg of total RNA from apices of D7 seedlings were separated on 17.5% PAGE-urea gels, blotted, and cross-linked to Hybond NX (Amersham ref. RPN203T) nylon membrane, as previously described ^65^. Probe hybridization was performed in PerfectHyb Plus buffer (Sigma ref. H7033) overnight at 42°C, followed by three 15-min washes in 2 x SSC 2% SDS at 48°C. MiRNA160 and U6 probes were obtained by labeling DNA oligonucleotides through PNK reaction with γ^32^ATP. To detect transposon-derived siRNA, PCR products were labeled with α^32^CTP through Klenow reaction. All primers and oligos for synthesis of probes are listed in Supplementary Table 3.

## Data availability

All sequencing data generated in this study are available at the Gene Expression Omnibus under accession number GSE192611

## Author contributions

G.B., V.H.N., O.M.S. and R.G conceived and designed the study and wrote the manuscript. G.B., V.H.N. and R.G. performed experiments and analyzed data. M.I. conducted northern blots, Z.M. supported snRNAseq analysis, H.B. made the CRISPR *ago4-CR* allele and supported sRNA analysis, M.D. made pElvis, N.L. supported several experiments. All authors read and approved the manuscript.

## Acknowledgments

We deeply thank the following colleagues: Matt Watson for manuscript revisions, Michael Borg for advice and support of pollen nuclei sorting, Michael Schon for initial support with snRNA-seq analysis, Alina Pröll, Fabia Kail, Johannes Rötzer for support with cloning, Thomas Lendl for support of microscope image analysis, Elin Axelsson-Ekker and Rahul Pisupati for helpful discussions and support with data analysis, Eriko Sasaki and Magnus Nordborg for helpful discussions.

We are grateful for excellent support by the GMI/IMP/IMBA Biooptics facilities, the Next Generation Sequencing and Plant Sciences units of the Vienna BioCenter Core Facilities (VBCF). We gratefully acknowledge financial support from the Austrian Science Fund (FWF I489, I1477 to O.M.S and I3687 to R.G.), Cost Action 16212 “INDEPTH” to O.M.S., and the Plant Fellows program (EU FP7) to R.G. We also acknowledge financial support for sequencing experiments via INTERREG RIAT-CZ to R.G.

