## Supplemental Material and Methods for "Two AGO proteins with transposon-derived sRNA cargo mark the germline in Arabidopsis"

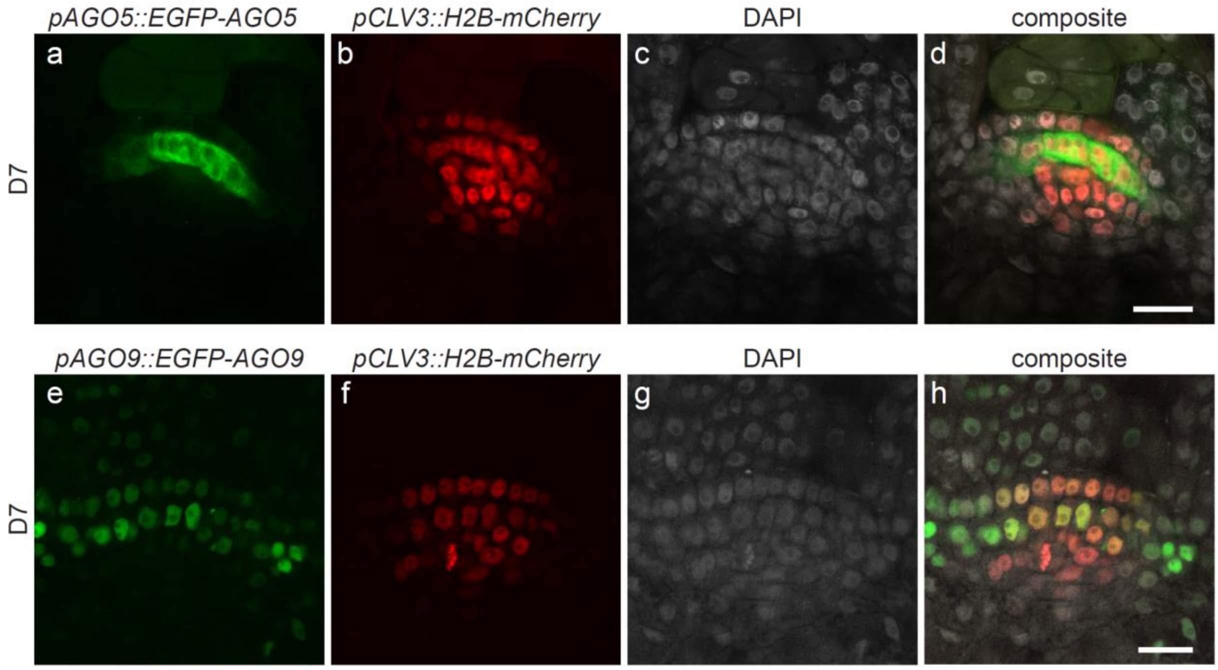


**Supplementary Fig. 1 | Localization of AGO5 and AGO9 in the SAM at D7.** Upper panel: localization of AGO5 in the SAM at D7, single channels and composite image. Lower panel: localization of AGO9 in the SAM at D7, single channels and composite image. (a, e) GFP; (b, f) mCherry; (c, g) DAPI; (d, h) composite image. Scale bar d, h = 20 µm. Images d and h are also used in Fig. 1a, b.

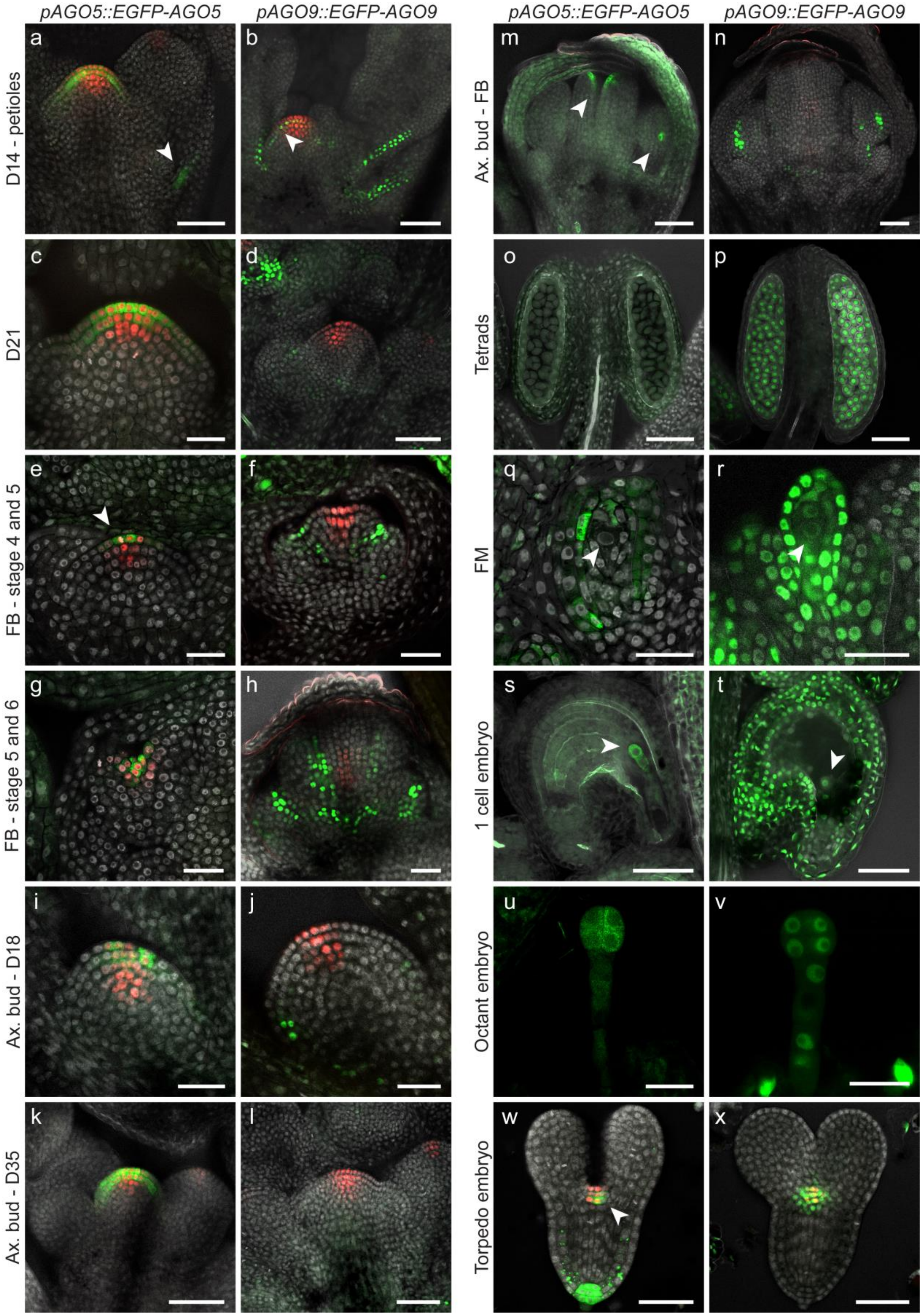


8

9 **Supplementary Figure 2 | Localization of AGO5 and AGO9 translational reporters at different**
10 **developmental stages. a**, AGO5 is localized in the L1, L2, and on the adaxial side at the base of leaf
11 petioles (arrowhead), where axillary meristems emerge. **b**, AGO9 is localized explicitly in the L2, and

the signal is found, as a continuum, along the adaxial side of leaf petioles. **c**, At D21, AGO5 maintains its localization in the two upper layers of the apical dome, and (**d**) AGO9 is not detected in the SAM anymore but present in the developing flower buds. **e**, In stage 4 flower buds, AGO5 is localized in the L1 of the whorl of carpels (arrowhead). **f**, At stage 5 of flower development, AGO9 is exclusively localized between the whorl of carpels and stamens. **g**, At Stage 5, AGO5 localization in the L1 follows the morphological changes that start shaping the two carpels and, at this point, it overlaps with stem cell nuclei. **h**, At Stage 6, AGO9-labeled nuclei span from the base of the flower to different cell layers in the upper developing organs. AGO9 is localized as a continuum between the carpel margin meristem (CMM) and the inner layers of the stamens. **i**, AGO5 is localized in the L1 and L2 of axillary buds at D18. **j**, AGO9 is localized at the periphery of axillary buds at D18. **k**, **l**, At D35, localization of AGO5 and AGO9 in meristems and **m**, **n**, flowers derived from elongated axillary buds resemble the pattern described in e-h. Arrowheads indicate the localization of AGO5. **o**, AGO5 is absent from microspore tetrads in contrast to **p**, AGO9. **q**, At Stage 4 of ovule development, AGO5 is restricted to the L1 and inner integuments and absent from the functional megaspore (FM) (arrowhead). **r**, AGO9 is localized to the FM (arrowhead) in the L1 and the surrounding integuments of Stage 4 ovules. **s**, AGO5, and (**t**) AGO9 are localized in the 1-cell embryo and the suspensor (arrowheads). **u**, AGO5, and (**v**) AGO9 are present in the body and suspensor of the octant stage embryo. **w**, Localization in the torpedo stage reveals the same pattern for AGO5 as observed in the heart stage (Fig. 1u) (arrowhead). **l**, In the torpedo stage, the AGO9 domain is broader than the CLV3 domain, and it reaches, to some extent, the basal regions of the cotyledons. Scale bar c, e, f, g, h, i, j, q, r, u, v = 20 µm. Scale bar a, b, d, k, l, m, n, o, p, s, t, w, x = 50 µm.

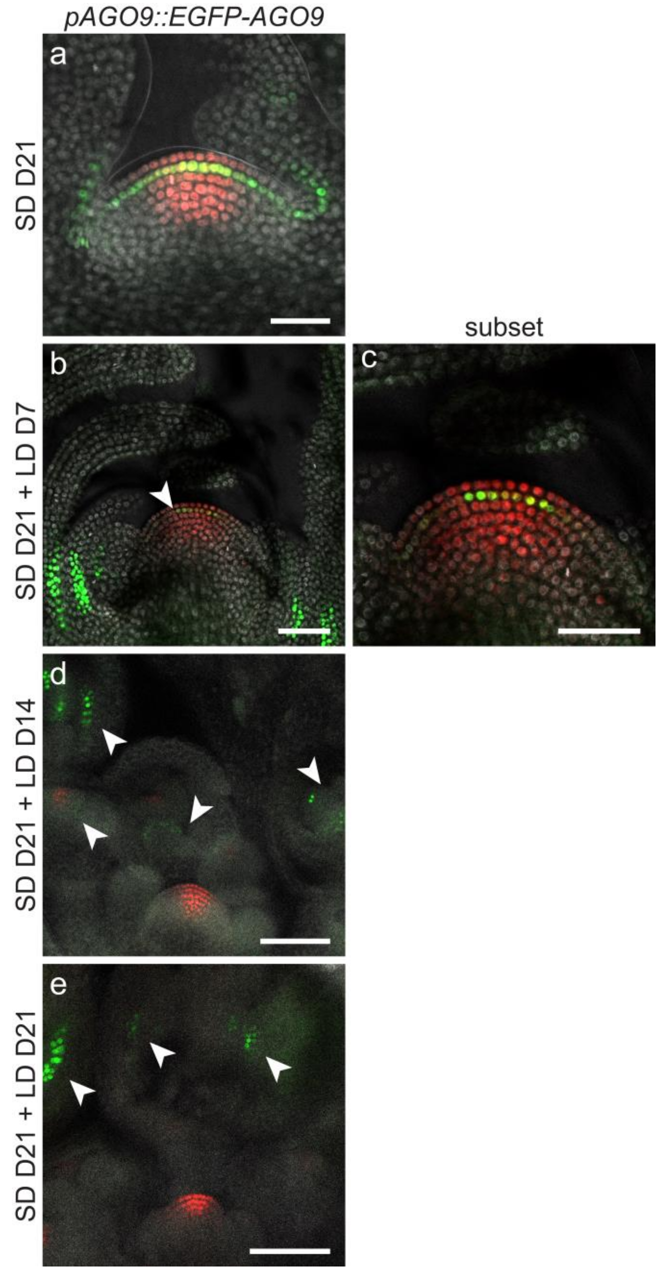


**Supplementary Figure 3 | Localization of AGO9 translational reporters during SD to LD transition. a**, At D21 in short days (SD), AGO9 is specifically localized in the L2 of the SAM and in leaf primordia**. b**, Three weeks in SD followed by a week of long days (LD D7) results in minor localization changes with AGO9 signal intensity only decreasing in intensity in the L2 (arrowhead) compared to (b)**. c**, After three weeks of SD and two (D14), or three (D21)(**d**) weeks of LD, AGO9 has disappeared from the SAM and can be detected in flowers instead (arrowheads). Scale bar a, b = 50µm. Scale bar c, d = 100µm.

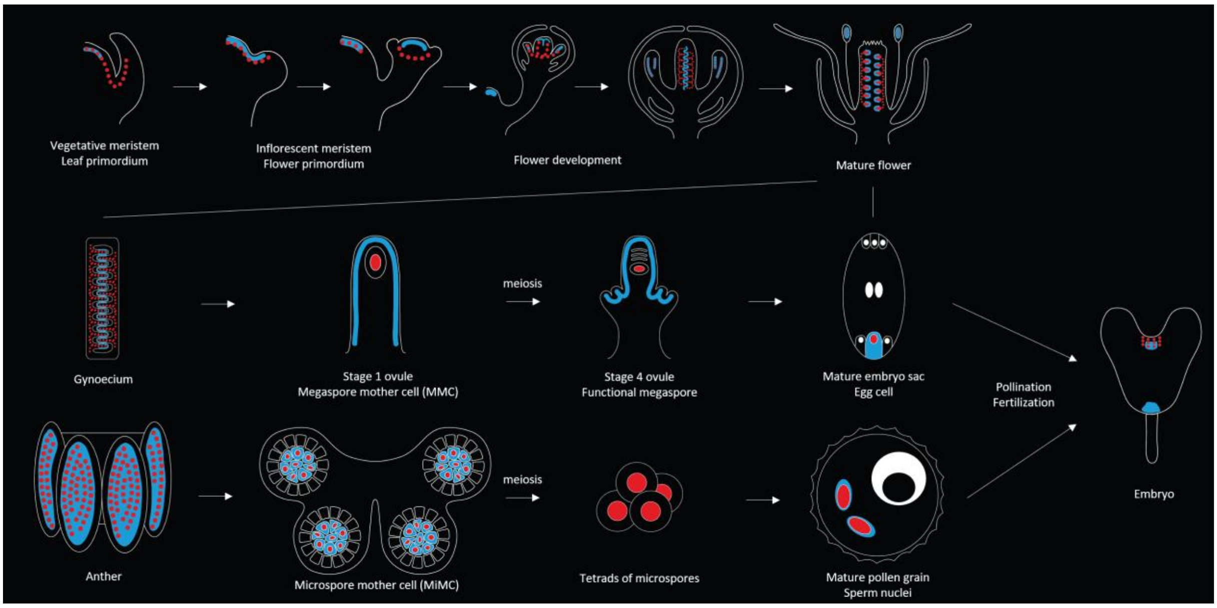


**Supplementary Fig. 4 | Schematic representation of AGO5 and AGO9 localization patterns throughout key stages of plant development.** AGO5-labelled cells are represented by continuous light blue lines. AGO9 labelled cells are represented by red dots.

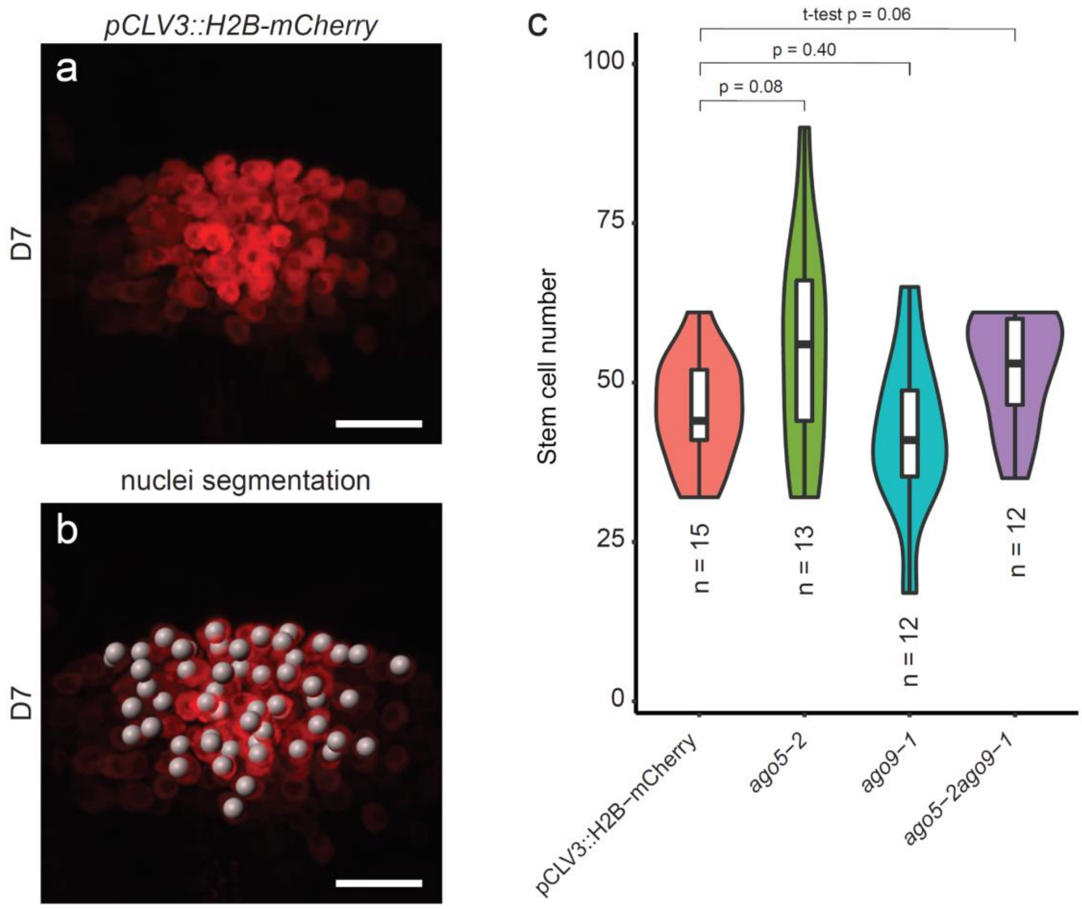


**Supplementary Fig. 5 | Number of *pCLV3::H2B-mCherry*-marked stem cells in D7 seedlings. a**, Representative image of *pCLV3::H2B-mCherry*-marked stem cells in a D7 seedling. **b**, Nuclei segmentation performed by Imaris software and overlaid on image in a. **c**, The number of stem cells in different lines. All mutant lines are in the *pCLV3::H2B-mCherry* background. n indicates the number of analyzed seedlings. Statistics were performed by means of t-test. Scale bar a, b = 10 µm.

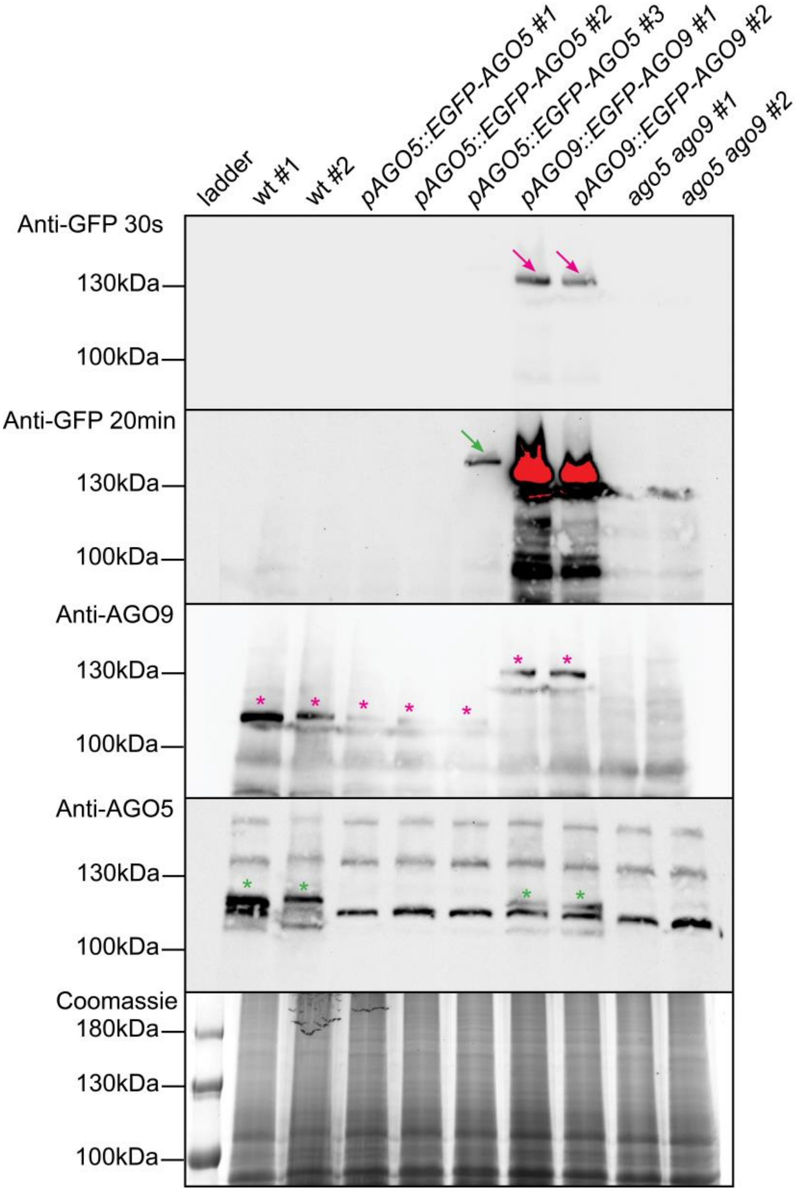


**Supplementary Fig. 6 | Protein levels in pAGO5::EGFP-AGO5 and pAGO9::EGFP-AGO9 lines.** Western blots prepared from 75 µg protein/lane in denaturing buffer. The protein extract was prepared from floral tissue. Purple arrows show the position of EGFP-AGO9 and purple stars that of untagged AGO9. The green arrow shows the EGFP-AGO5 signal and green stars that of untagged AGO5. Only one of the tested pAGO5::EGFP-AGO5 lines has a detectable expression level and was used for the immunoprecipitation. The AGO5 antibodies do not detect EGFP-AGO5**.**

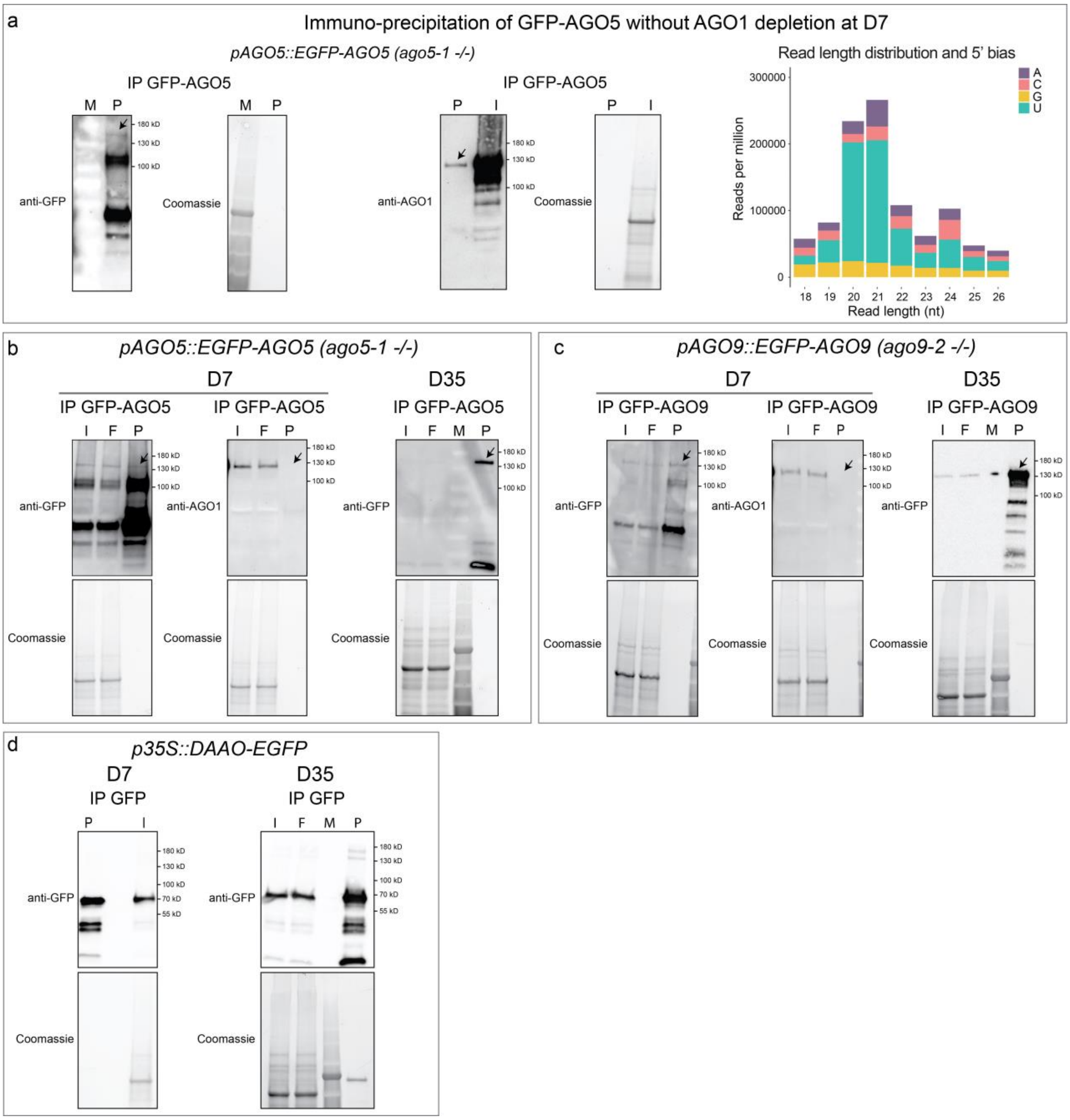


**Supplementary Fig. 7 | Immuno-precipitation of GFP-AGO5 and GFP-AGO9 at D7 and D35. a**, Initial analysis of AGO5-bound sRNAs at D7 revealed high content of 21 nt sRNAs with 5' U similar to sRNAs described associated with AGO1 (shown on the right side). Therefore, we tested the AGO5 precipitate for AGO1 contamination. Indeed, AGO1 was detectable in the AGO5 precipitate (arrow, Western in the middle). **b**,**c**,**d**, Prior to AGO5 or AGO9 precipitation, we included an AGO1 depletion step and controlled for the absence of AGO1 in the precipitate. Immunoprecipitation of GFP-AGO5 (b), GFP-AGO9 (c), and GFP control (d) at D7 and D35. For the GFP-control (lower panel) we used a line expressing a constitutive p35S::DAAO-GFP. Arrows indicate corresponding signals. Nomenclature: IP (immuno-precipitation), P (precipitate), I (Input), F (Flow), M (protein ladder).

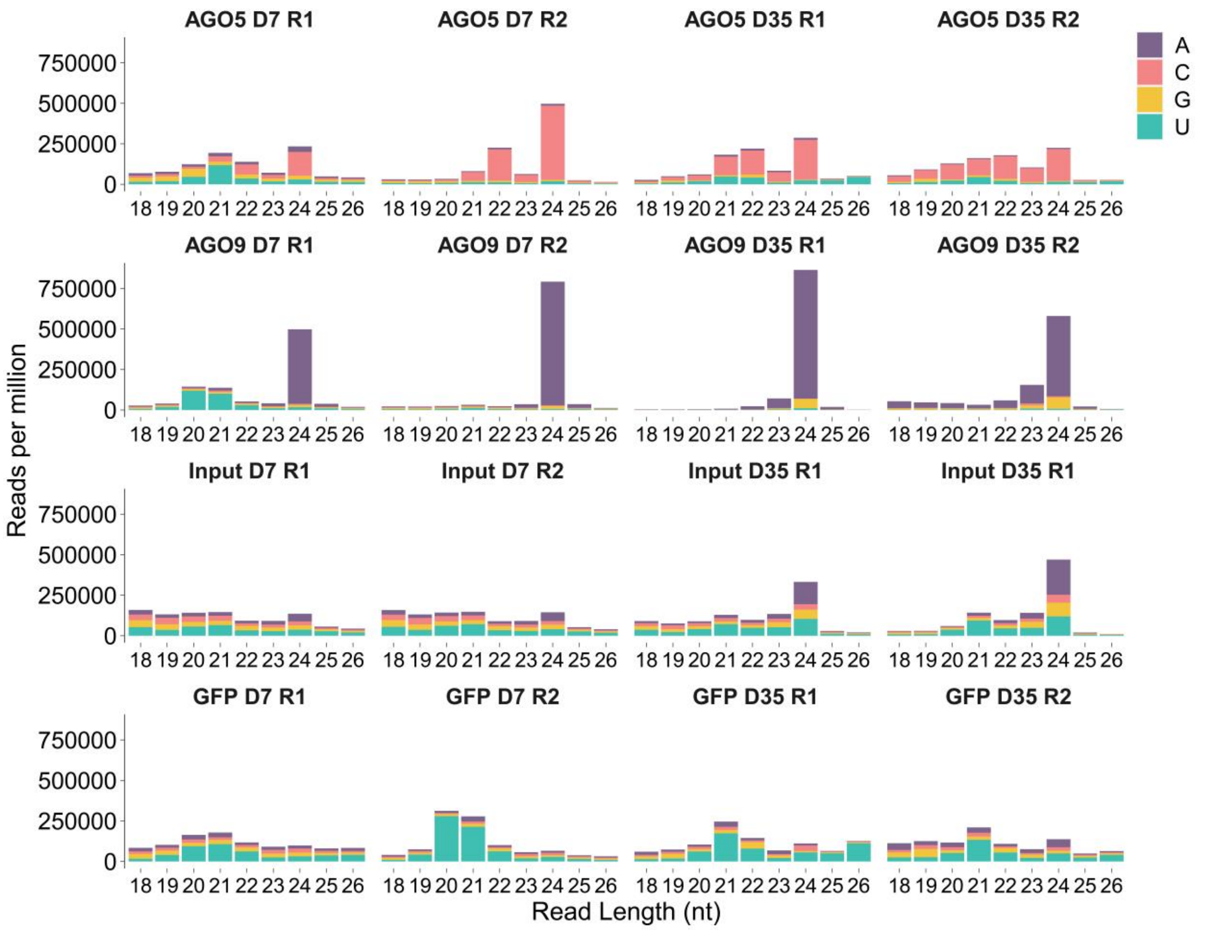


**Supplementary Fig. 8 | Read length distribution and 5' bias of AGO5- and AGO9-associated sRNAs**. Results are from two independent replicas (R1 and R2). For the GFP-control (lower panel), we used a line expressing a constitutive p35S::DAAO-GFP.

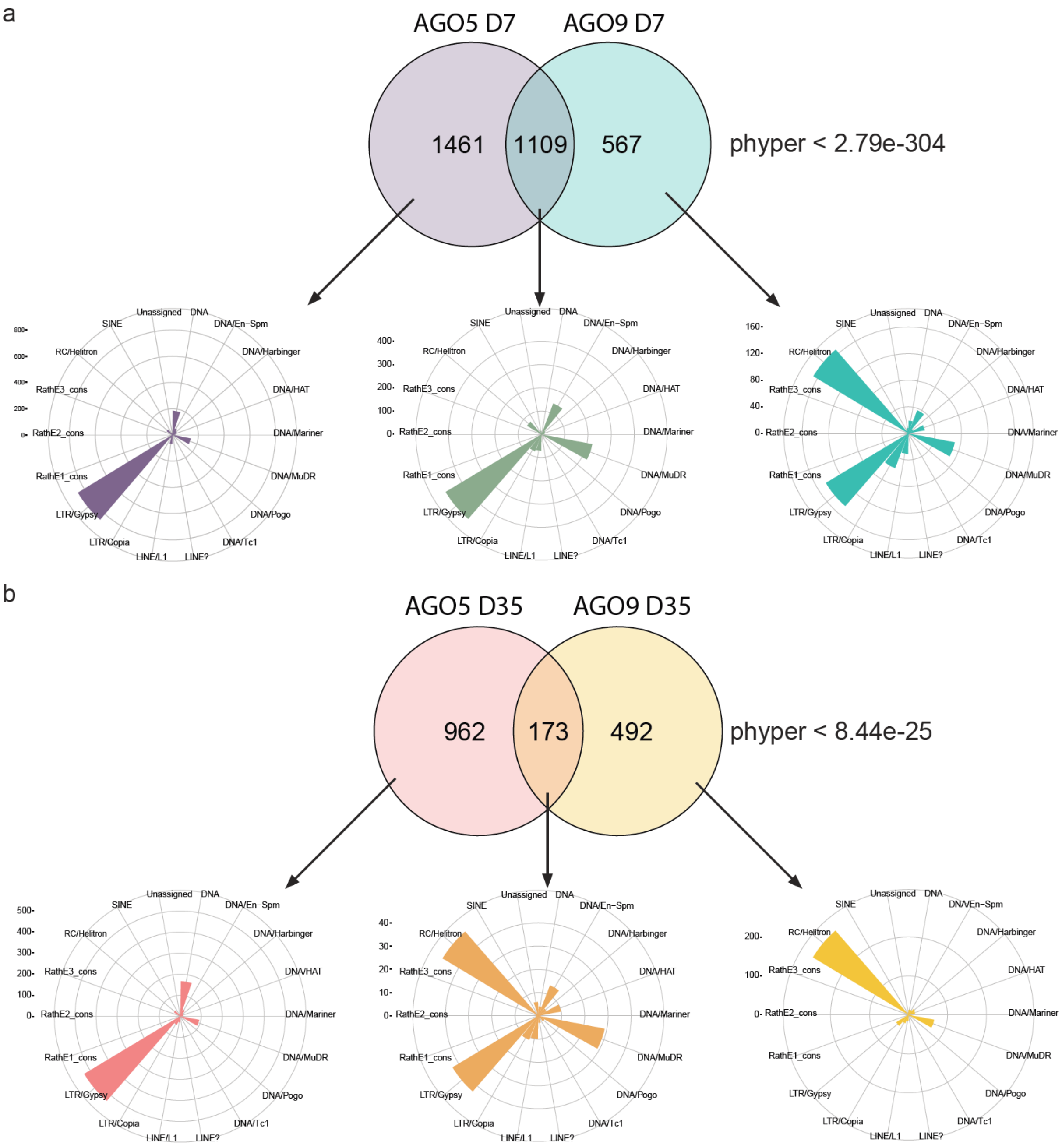


**Supplementary Fig. 9 | AGO5 and AGO9 cargo derives from common and specific transposons throughout development**. Venn diagrams show the number of overlapping targets and p-values at D7 (a) and D35 (b). The polar charts indicate composition of corresponding superfamilies.

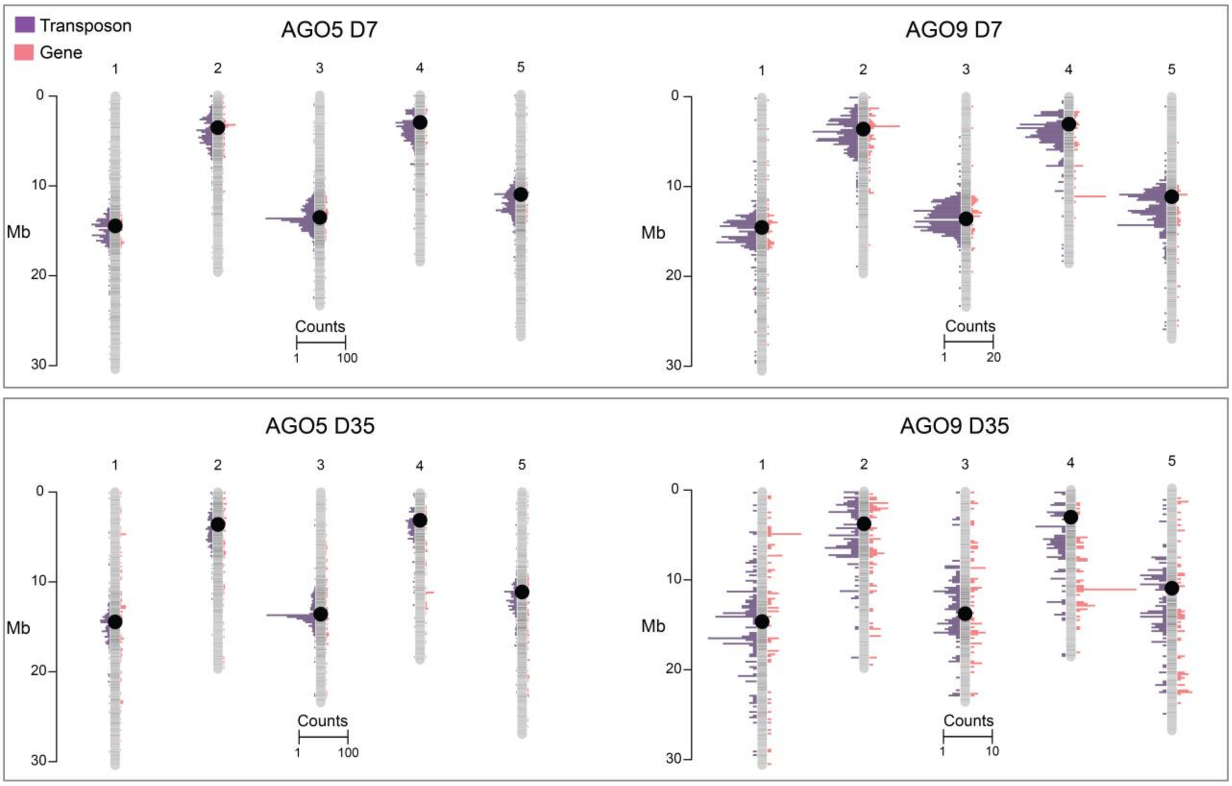


**Supplementary Fig. 10 | Annotation of AGO5- and AGO9-enriched cargo on the five chromosomes**. AGO5-bound sRNAs at D7 (a) and at D35 (c) and AGO9 bound sRNAs at D7 (b) and at D35 (d).

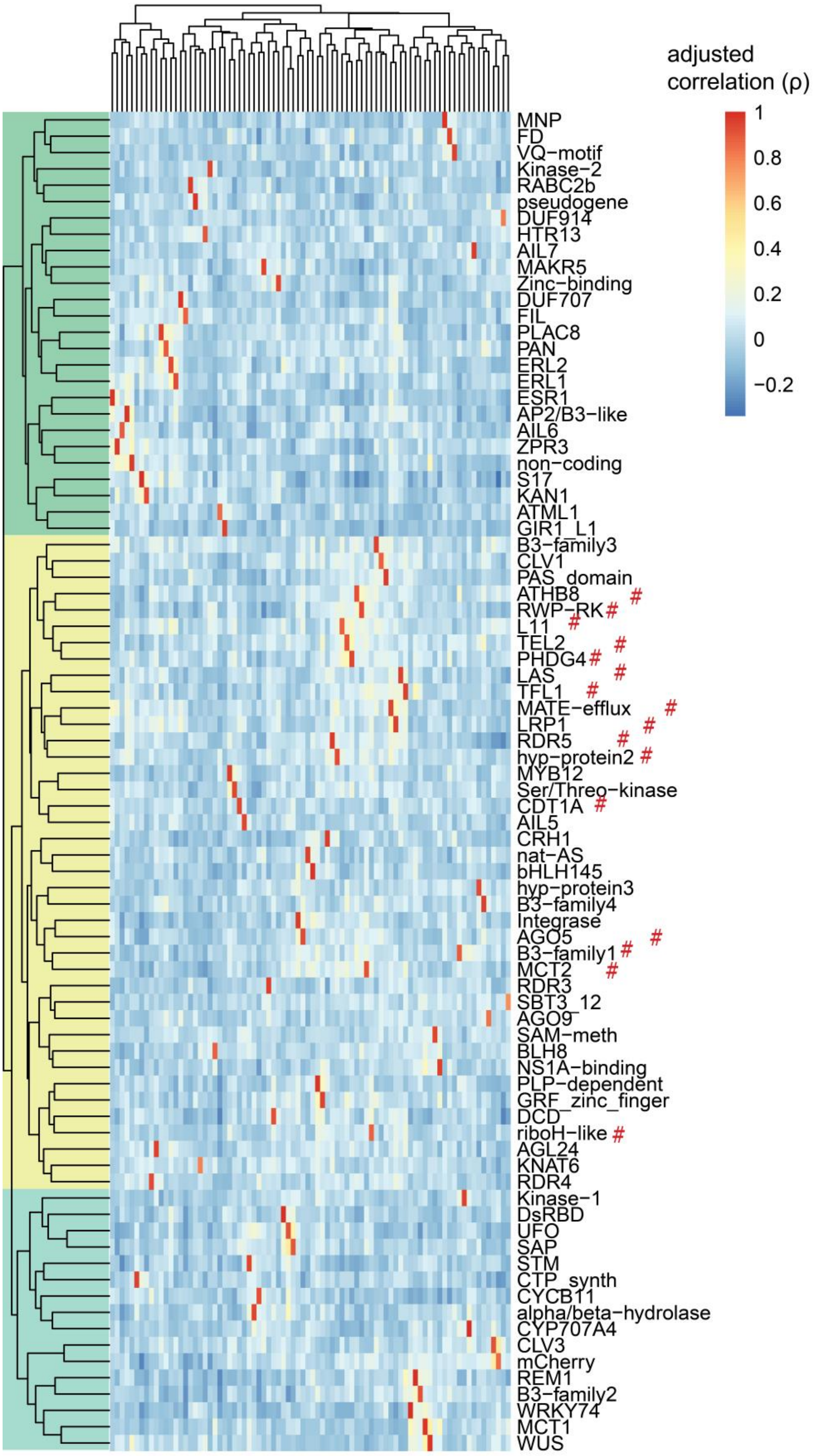


**Supplementary Fig. 11 | Clustered adjusted correlation matrix of GESS.** Shown is an adjusted correlation matrix for 79 GESS and 3 cell cycle reporters (82X82 gene-gene correlations). Genes marked with # cluster together in a genome-wide correlation matrix as shown in Fig. 4.

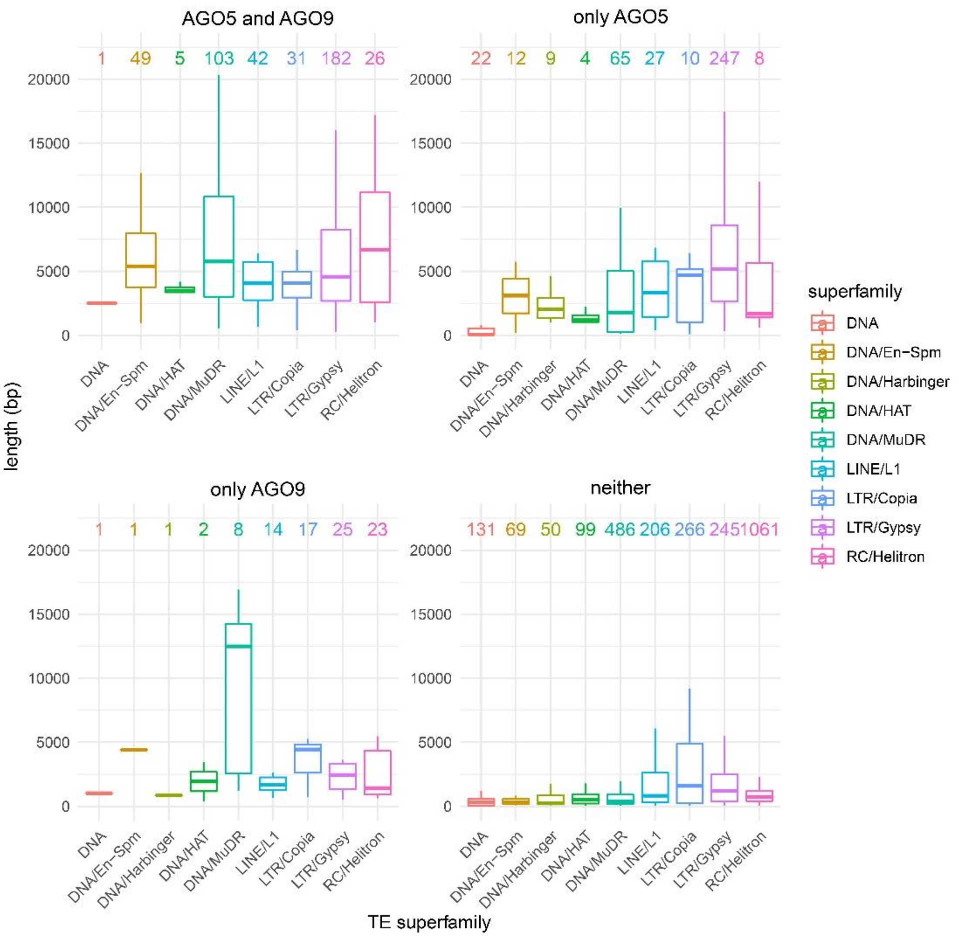


**Supplementary Fig. 12 | Length of TEs expressed in single SAM stem cell nuclei.** Groups indicate whether TEs are processed into sRNAs associated with AGO5 and AGO9, only AGO5, only AGO9, or neither. Coloured numbers show number of TEs in the respective superfamily.

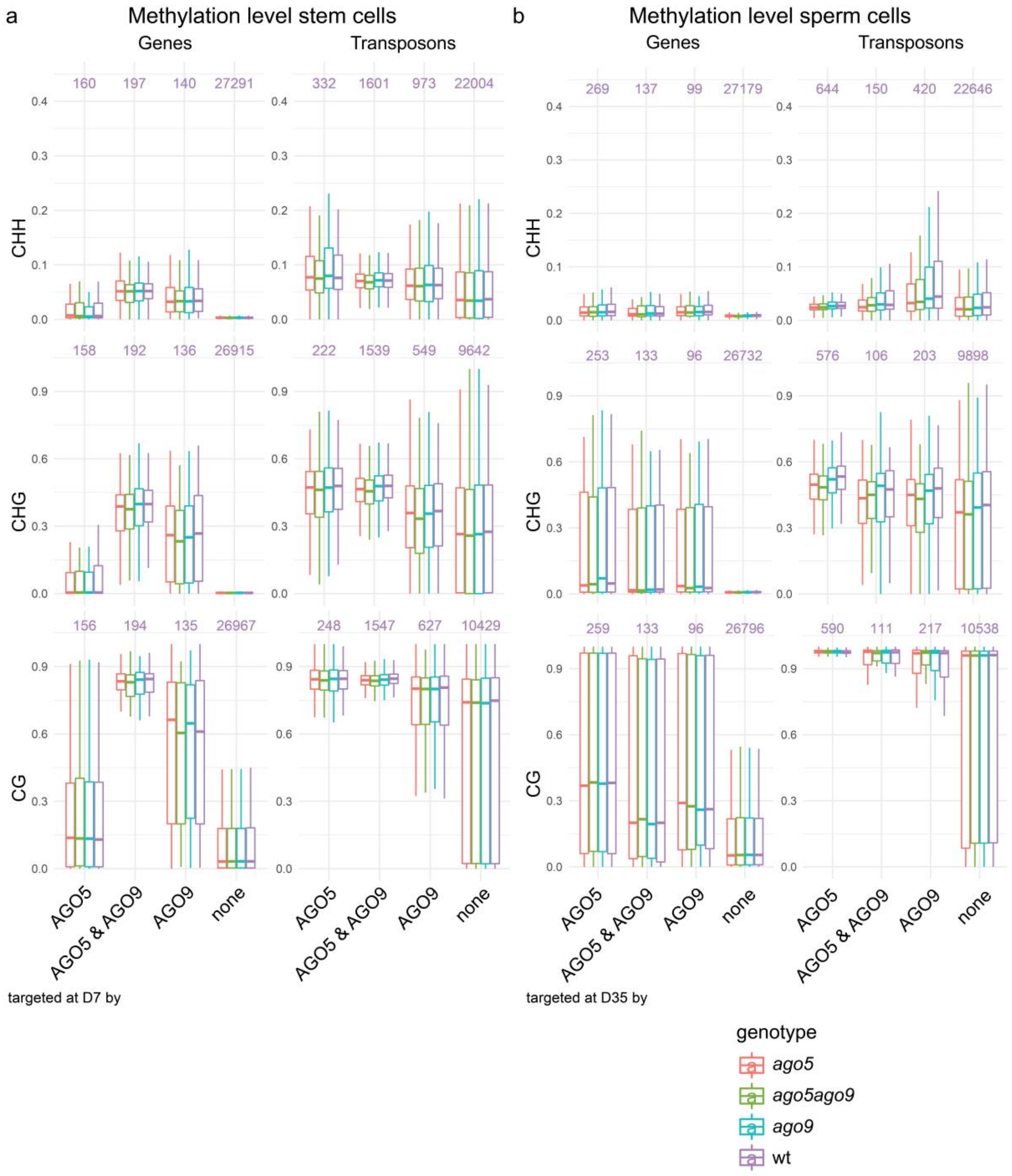


**Supplementary Fig. 13 | Methylation levels on genes and TEs. a**, Methylation levels of stem cells of *ago5*, *ago9*, *ago5 ago9*, and wt on genes and TEs targeted either by AGO5, AGO9, both, or none, at D7. Purple numbers indicate number of targets. **b**, Methylation levels of sperm cells of *ago5*, *ago9*, *ago5 ago9*, and wt, on genes and TEs targeted either by AGO5, AGO9, both, or none, at D35. Purple numbers indicate number of targets.

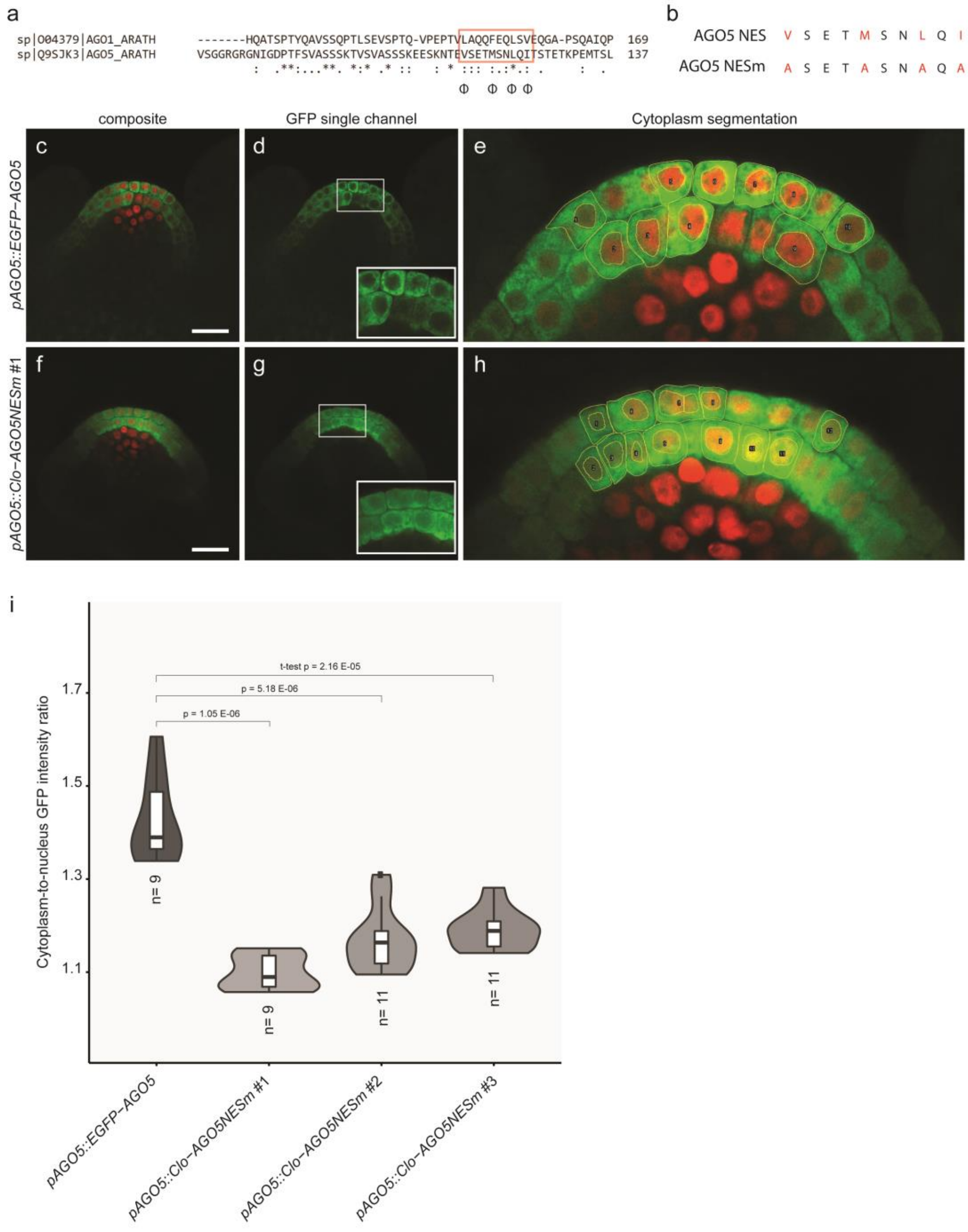


**Supplementary Figure 14 | Mutation of the AGO5 nuclear export signal leads to nuclear accumulation of AGO5. a**, Clustal O (1.2.4) amino-acid sequence alignment between AGO1 and AGO5. The red box identifies the presumptive NES, and Φ represents either L, I, V, M, or F, as described for

AGO1 (Bologna et al. 2018). **b**, Alanine substitutions are indicated for AGO5 NESm. **c**, Representative image of *pAGO5::EGFP-AGO5* at D35. **f**, Representative image of *pAGO5::Clo-AGO5NESm* #1 at D35. **d**, **g**, GFP single channel of image c and f; insets: apical GFP labelled cells. **e**, **h**, Subset of output image derived from automated cytoplasm segmentation of image c and f. Yellow lines outline the nucleus and cytoplasm perimeter of cells. **i**, Cytoplasm-to-nucleus GFP intensity ratio of L1 and L2 apical cells in D35 plants. All lines are in the *pCLV3::H2B-mCherry* and *ago5-1* mutant background. n indicates the number of apices analyzed. Between 9 to 21 cells per apex were used for the calculations. Statistics were performed employing a t-test. Scale bar c and f = 20 µm.

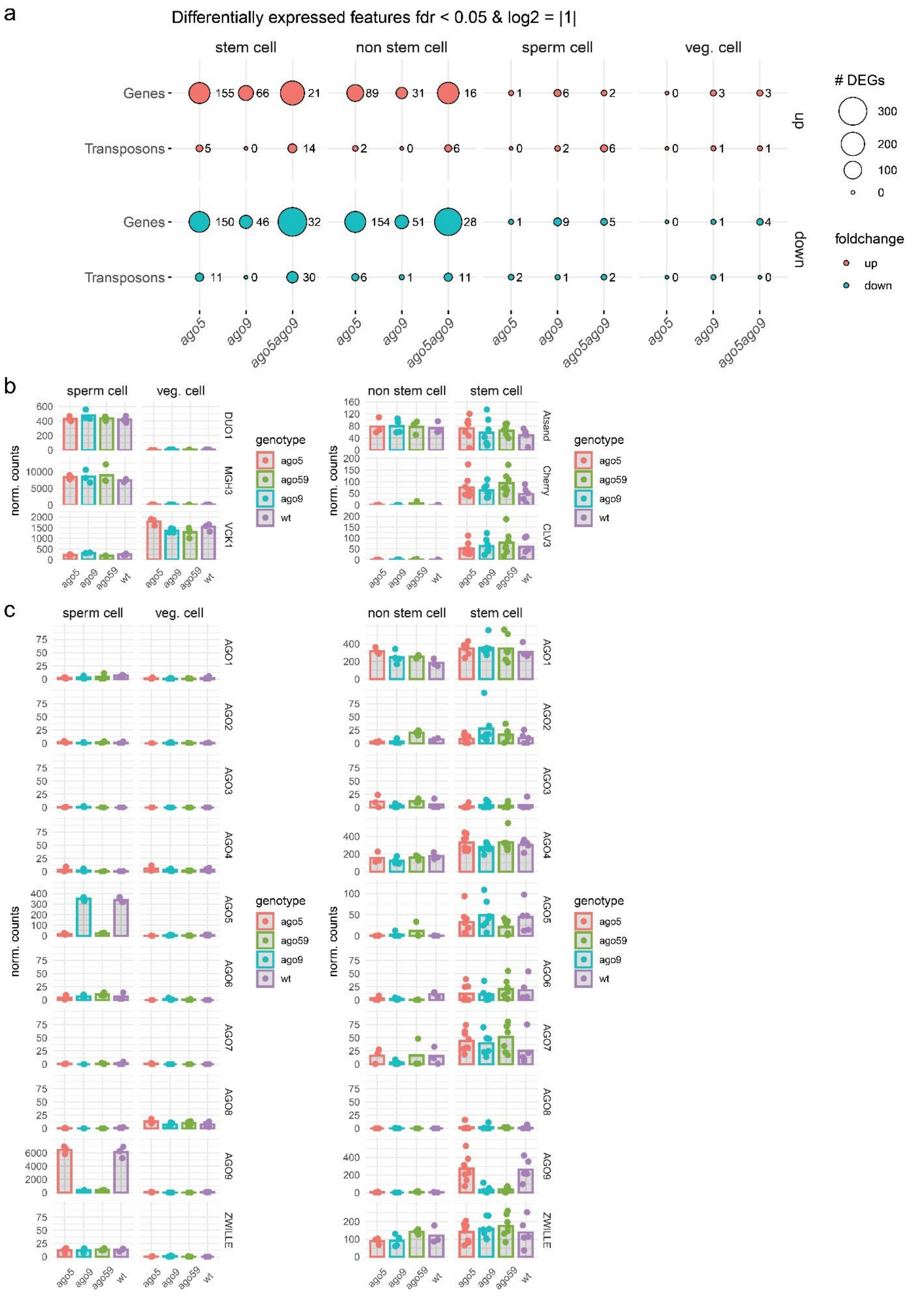


**Supplementary Fig. 15 | mRNAseq analysis of sorted stem, non-stem, sperm, and vegetative (vn) nuclei. a**, Balloon plot representing the number of DEGs for comparison with the wt. **b**, Expression of *DUO1*, *MGH3* (reporter for sperm cells), *VCK1* (reporter for vn cells), *CLV3*, mCherry (reporter for stem cells), and *AtSand* (ubiquitously expressed). **c**, Expression of all 10 *AGO* genes. Reads aligning to *AGO5* in stem cells of *ago5* and *ago5 ago9* (ago59) map to the region before the T-DNA insertion in the mutant.

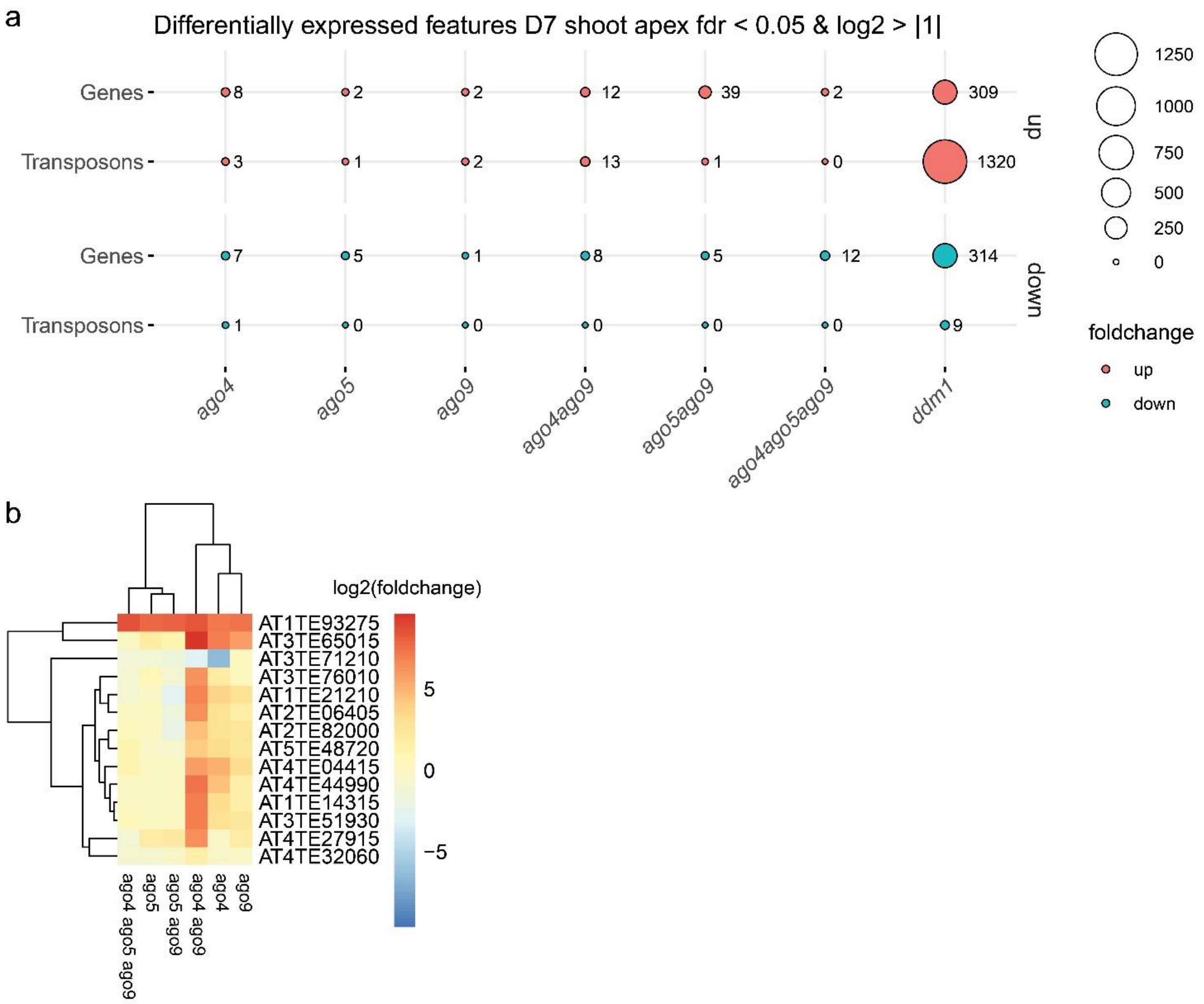


**Supplementary Fig. 16 | mRNAseq analysis of D7 shoot apices. a**, Number of differentially expressed genes and TEs (DEGs) in D7 shoot apices of indicated mutants compared to wild type. **b**, Clustering of log2-fold changes of TEs with increased expression in *ago* mutants showing synergistic effects of AGO4 and AGO9 on the expression of transposons. Some differences between *ago4 ago9* and *ago4 ago5 ago9* might originate from different mutant alleles for *ago4*, as we had to generate the allele in the triple mutant with CRISPR due to the genetic linkage of *AGO4* and *AGO5* on chromosome two.

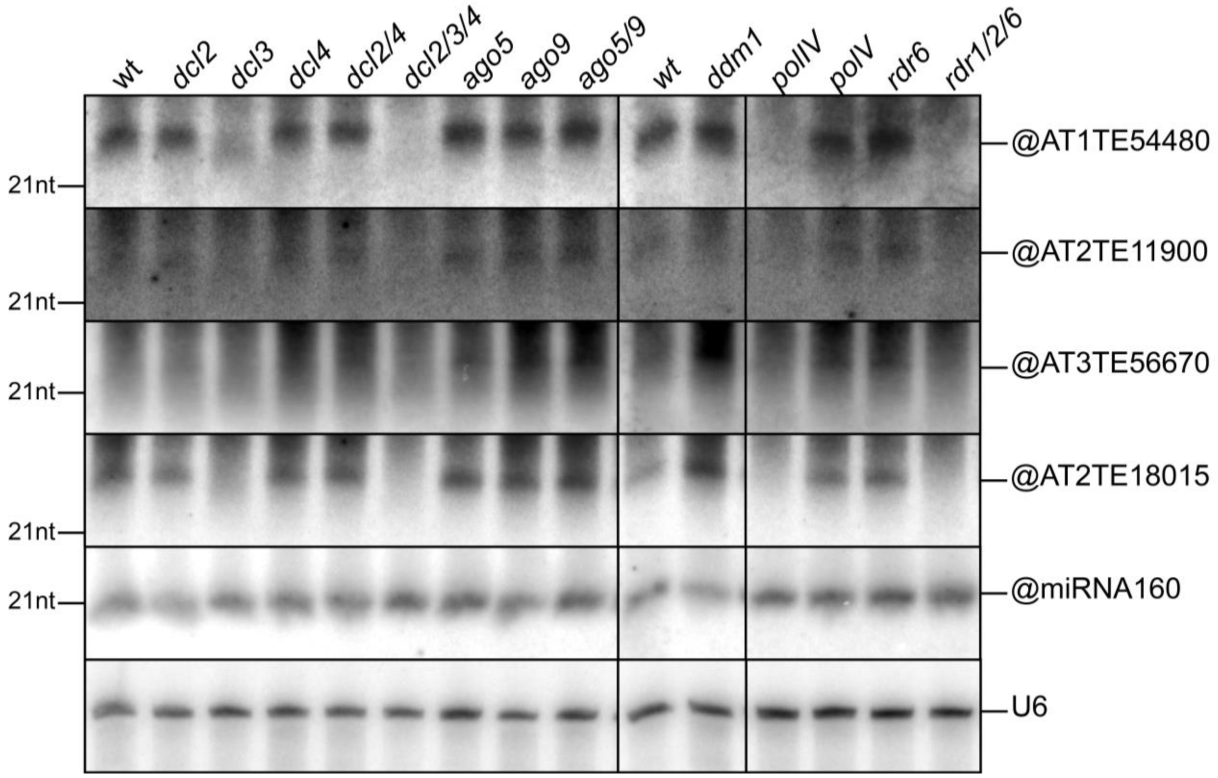


**Supplementary Fig. 17 | siRNAs derived from TEs targeted by AGO5 are synthesized by PolIV, RDR2, and DCL3.** RNA was extracted from apices of 7 day-old seedlings of the indicated genotypes. sRNAs from the indicated TEs show no or a weak signal in *dcl2*, *dcl2/3/4*, *polIV*, and *rdr1/2/6* mutants, but a visible signal in the size class from 22-24 nt in the other genotypes. miRNA160 and U6 are shown as loading controls.

### Supplementary Methods

#### Alignment and counting of transposable elements.

With the release of the Arabidopsis genome annotation Tair8, a new transposon annotation, based on multiple homology-based predictions, has been added [1]. Existing annotations, overlapping with TE annotations, have been reclassified as locus type “transposable element gene” (https://arabidopsis.org/download_files/Genes/TAIR8_genome_release/Readme-transposons). For alignment and assigning sequencing reads to either genes, transposons (TEs), or transposable element genes (TE genes), this creates a problem of redundant annotations. Supplementary Fig. 18 shows a large proportion of TE genes overlapping with more than one TEs and vice versa. For avoiding assigning reads to both overlapping TE genes and TEs, we removed TE genes from the Tair10 annotation and added TEs as a single feature type (Tair10+TEs).

Alignment strategies can vary in their accuracy and resolution, especially for TEs [2,3]. To find an optimal alignment and feature counting method, we compared several alignment and quantification tools with a test data set and used DESeq2 for calculating differentially expressed features (genes and transposons). We proceeded using STAR for alignment and Salmon for read quantification, as this resulted in the smallest number of private DEGs (Supplementary Fig. 19).

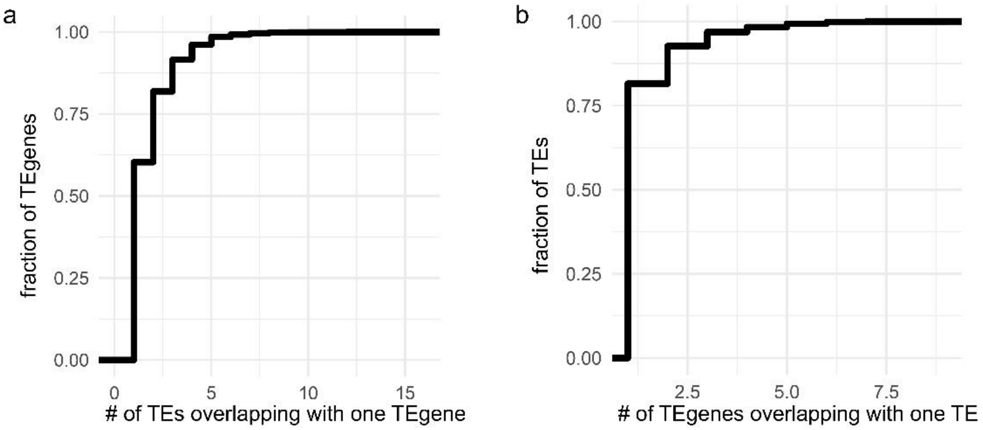


**Supplementary Fig. 18 |** Fraction of TE genes with more than one overlapping TEs (**a**) and the fraction of TEs with more than one overlapping TE genes (**b**).

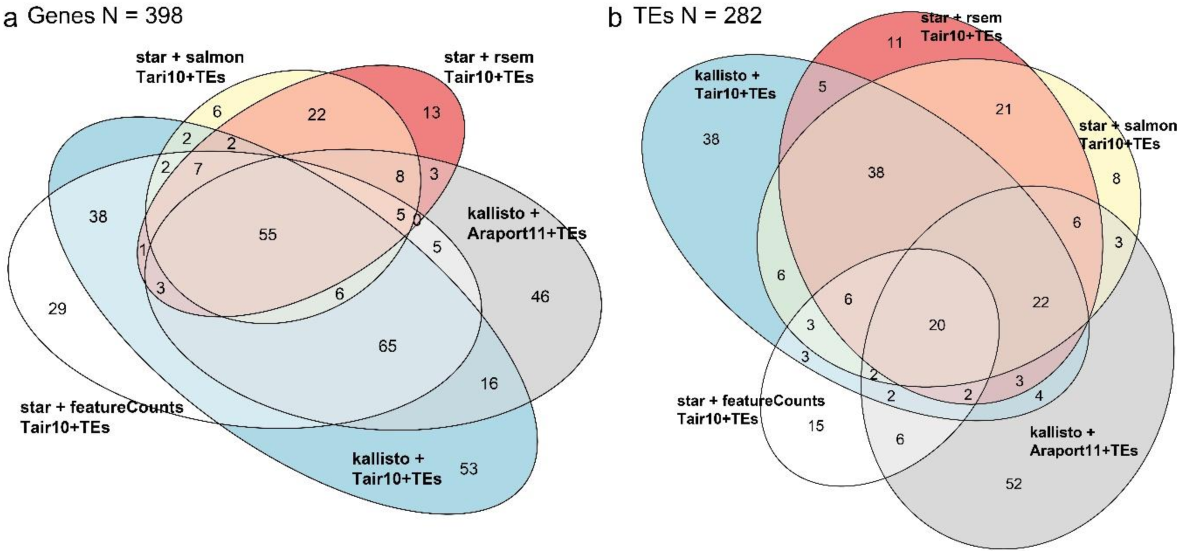


**Supplementary Fig. 19 |** An example dataset was aligned with either star or kallisto and reads quantified with featureCounts, Salmon, or Kallisto. Shown is the overlap of DESeq2 differentially expressed features (FDR < 0.05) for each count table for genes (**a**) and transposons (**b**).

**Analysis of single nuclei sequencing data.**

Before sorting single stem cell nuclei, sorting accuracy was confirmed using counting of nuclei (by microscopy) and quantitative PCR for mRNA and genomic DNA. For sequencing, a total of 208 single nuclei were sorted into 96-well plates. We included two bulk controls of 50 nuclei and two empty negative controls. A count matrix was generated as described above (using STAR for alignment and Salmon for read quantification). 20 nuclei with low feature and read count were filtered out, resulting in the feature count distribution of Supplementary Fig. 20a. 217 genes with very high read counts and variance and mostly encoding genes for translational or photosynthetic processes were filtered out. We chose four as a cut-off based on the number of features expressed in a certain number of nuclei (Supplementary Fig. 20b). Therefore, each feature (gene or transposon) was expressed in at least four nuclei. We performed an index sort for one plate and recorded every nucleus's DAPI and mCherry intensities. Sorting order did not correlate with the number of detected features, showing that mRNA leakage of nuclei during sorting is not problematic (Supplementary Fig. 21a). Surprisingly, DAPI, but not mCherry intensities, were highly correlated with the number of detected features (Supplementary Fig. 21b-d). This shows that the cell cycle state of the nuclei contributes strongly to variation in the number of detected genes. This correlation was even slightly higher than the correlation of the number of detected genes with the number of aligned reads. We also could assign a cell cycle state to more than 90 nuclei based on the expression of *HTR13* (S-G2), *CDT1A* (G1), and *CYCB1.1* (G2-M) (Supplementary Fig. 22). For calculating and clustering gene-gene correlations, we first computed Spearman's correlation between all genes, then adjusted the correlation value between every pair of genes by their sampling depth [4]. In brief, this strategy subtracts the expected correlation between any pair of genes based on their expression levels only. This allows to detect notable correlations between genes even if they were lowly sampled (which is typically the case in sparse datasets such as snRNAseq) and vice versa, to not overestimate high correlation values between well-covered genes. As an example, we show correlation and adjusted correlation values for *CLV3* (Supplementary Fig. 23). For calculating TE abundance in different nucleus-types, data were processed with DESeq2.

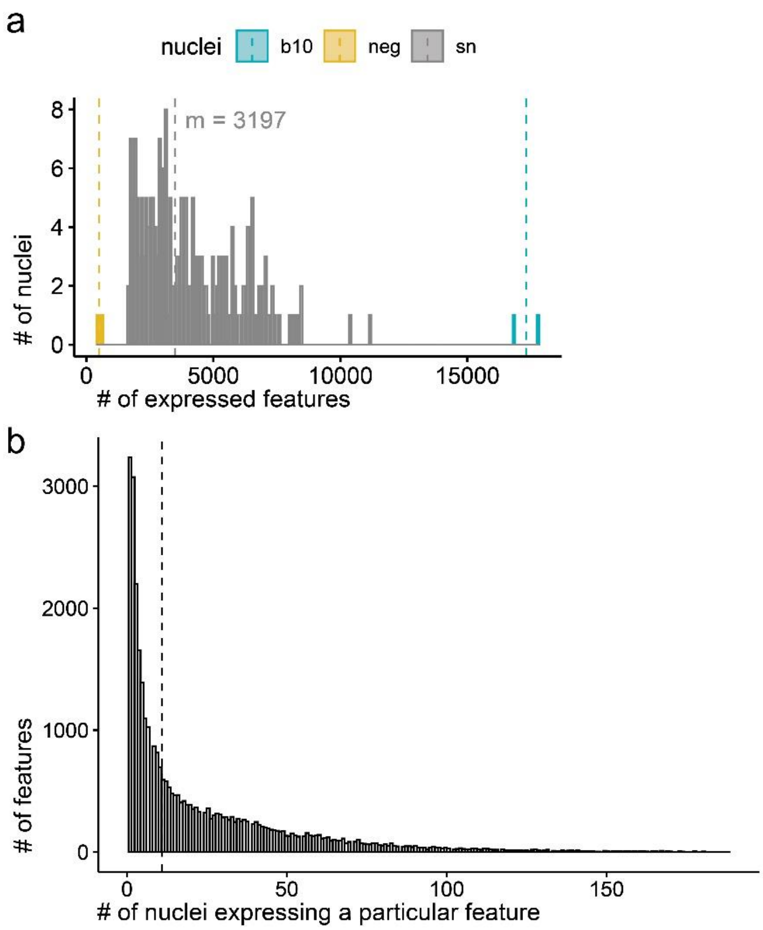

**Supplementary Fig. 20 | a**, Number of detected features (genes and TEs) in single nuclei (sn), negative controls (neg), and bulk controls of 50 nuclei (b10). **b**, Number of features expressed in number of nuclei.

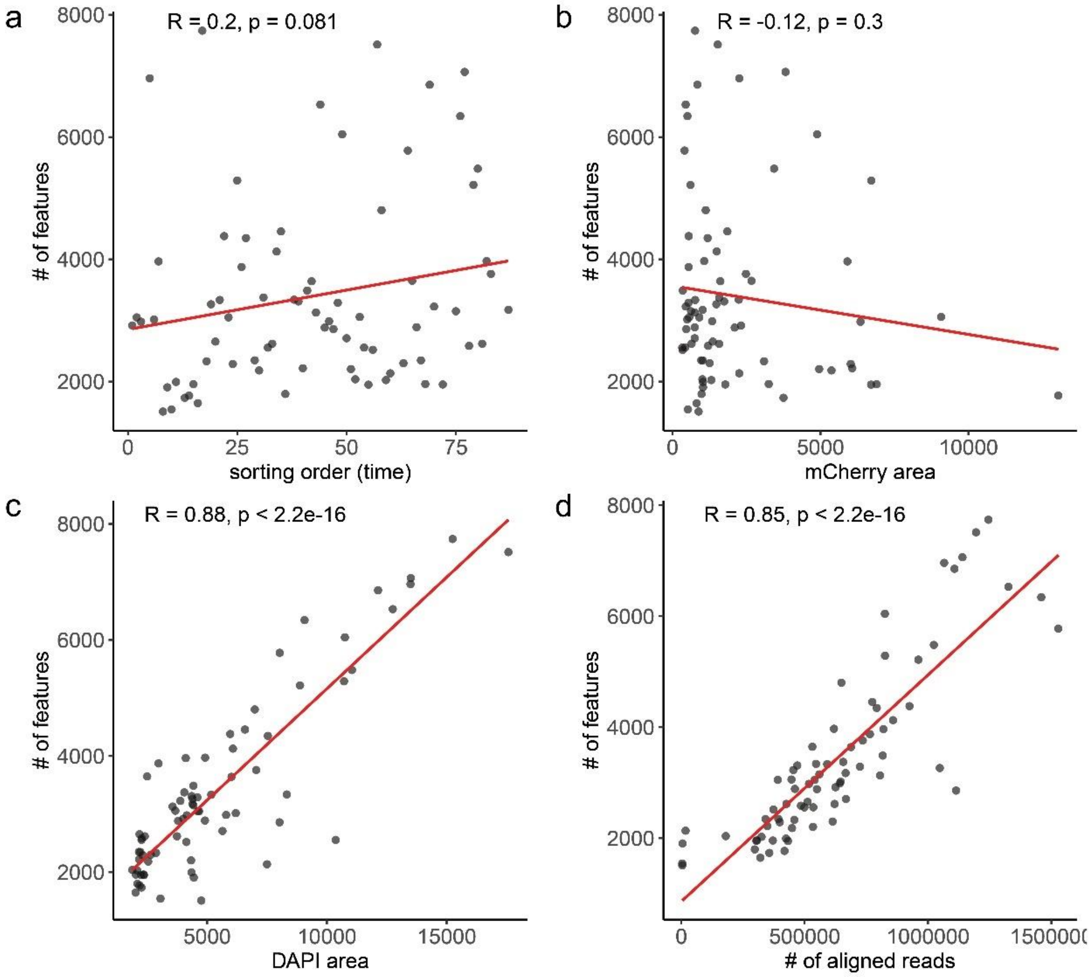


**Supplementary Fig. 21 | a,** Scatterplots showing # of detected features (genes and TEs) over sorting order, **b,** mCherry signal, **c,** DAPI signal, and **d,** number of aligned reads.

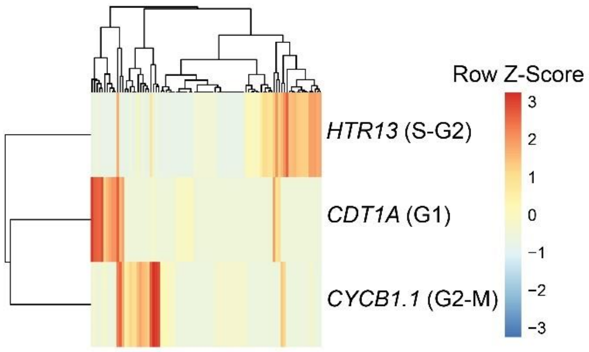


**Supplementary Fig. 22 |** Cell cycle state of individual SAM nuclei. Shown are expression values. Three cell cycle reporter genes separate individual nuclei into three major clusters.

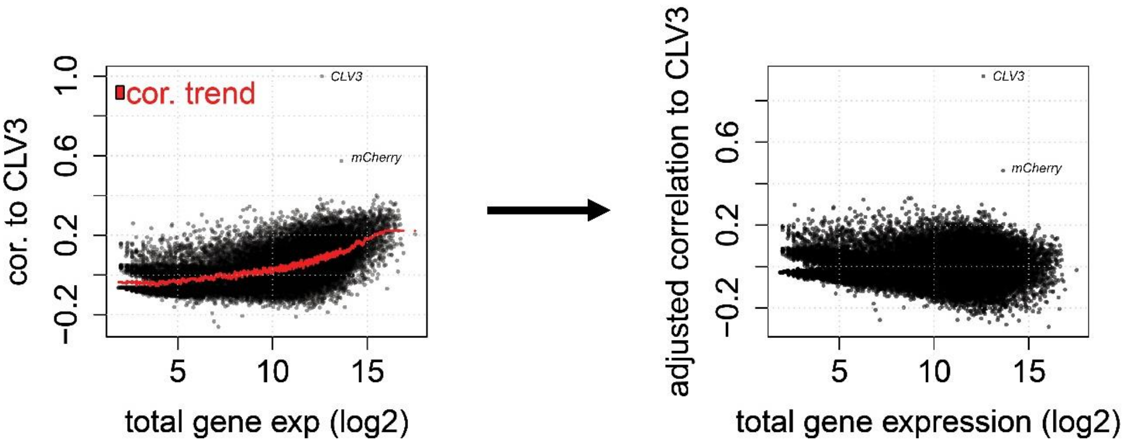


**Supplementary Fig. 23 |** Correlation and adjusted correlation to *CLV3*. Each dot resembles a gene (N= 24,761) and its raw Spearman's correlation (left) and adjusted correlation (right) to *CLV3* are depicted on the y-axis. Total expression levels (log2 transformed) are shown on x-axis.

**References supplementary methods**


1. Buisine, N., Quesneville, H. & Colot, V. Improved detection and annotation of transposable elements in sequenced genomes using multiple reference sequence sets. *Genomics* **91**, 467-75 (2008).
2. Lanciano, S. & Cristofari, G. Measuring and interpreting transposable element expression. *Nat Rev Genet* **21**, 721-736 (2020).
3. O'Neill, K., Brocks, D. & Hammell, M.G. Mobile genomics: tools and techniques for tackling transposons. *Philos Trans R Soc Lond B Biol Sci* **375**, 20190345 (2020).
4. Meir, Z., Mukamel, Z., Chomsky, E., Lifshitz, A. & Tanay, A. Single-cell analysis of clonal maintenance of transcriptional and epigenetic states in cancer cells. *Nat Genet* **52**, 709-718 (2020).
